# Intraspecific genomic variation and local adaptation in a young hybrid species

**DOI:** 10.1101/732313

**Authors:** Angélica Cuevas, Mark Ravinet, Glenn-Peter Sætre, Fabrice Eroukhmanoff

## Abstract

Hybridization increases genetic variation, hence hybrid species may have a strong evolutionary potential once their admixed genomes have stabilized and incompatibilities have been purged. Yet, little is known about how such hybrid lineages evolve at the genomic level following their formation, in particular the characteristics of their adaptive potential, i.e. constraints and facilitations of diversification. Here we investigate how the Italian sparrow (*Passer italiae*), a homoploid hybrid species, has evolved and locally adapted to its variable environment. Using restriction site-associated DNA sequencing (RAD-seq) on several populations across the Italian peninsula, we evaluate how genomic constraints and novel genetic variation have influenced population divergence and adaptation. We show that population divergence within this hybrid species has evolved in response to climatic variation. As in non-hybrid species, climatic differences may even reduce gene flow between populations, suggesting ongoing local adaptation. We report outlier genes associated with adaptation to climatic variation, known to be involved in beak morphology in other species. Most of the strongly divergent loci among Italian sparrow populations seem not to be differentiated between its parent species, the house and Spanish sparrow. Within the parental species, population divergence has occurred mostly in loci where different alleles segregate in the parent species, unlike in the hybrid, suggesting that novel combinations of parental alleles in the hybrid have not necessarily enhanced its evolutionary potential. Rather, our study suggests that constraints linked to incompatibilities may have restricted the evolution of this admixed genome, both during and after hybrid species formation.

## INTRODUCTION

Hybridization is an evolutionary process that has been increasingly studied in the last decade (Abbott et al., 2013; Marques, Meier & Seehausen, 2019; Taylor & Larson, 2019). It can have a wide array of consequences, ranging from speciation reversal, reinforcement of prezygotic barriers to gene exchange, adaptive introgression and hybrid speciation. In particular, hybrid speciation – the formation of new species as a result of hybridization (Mallet, 2007) – can be seen as one of the most creative outcomes of hybridization. This is especially true in the case of homoploid hybrid speciation (HHS), which is thought to be rare given that reproductive isolation from the parental species does not automatically derive from differences in ploidy levels. Nevertheless, in the last decade, several compelling cases of HHS have been described in animals (Abbott et al., 2013; Mallet, 2007; Schumer, Rosenthal & Andolfatto, 2014). Most of these studies focused on making a case for demonstrating HHS while little focus has been put on analyzing the evolutionary fate and adaptive potential of hybrid species. In the long term, the establishment and success of a homoploid hybrid species only depend partially on the fast evolution of reproductive barriers that isolate it from its parental species and the purging of incompatibilities. Selection should also favor locally adapted allelic combinations to ensure the hybrid’s ecological persistence and further adaptation to its potentially variable environment.

Genetic variability in hybrid lineages can be produced by the admixture process itself, through the generation of heterozygocity at loci that are differentially fixed in the parental species, novel re-arrangements of parental ancestry blocks or the inheritance of parental standing genetic variation (i.e. loci where different alleles are still segregating in one or both of the parental species) (Abbott et al., 2013). These processes can induce a substantial amount of genetic variation in the hybrid lineage, which may later display a higher evolutionary potential than that found in non-hybrid species. Empirical studies have shown that novel genetic combinations in hybrid lineages can substantially increase phenotypic variation (Meier et al., 2017) and even lead to adaptive radiations (Keller et al., 2013). However, the evolutionary potential of a hybrid species can be hampered by the limitations generated by genomic incompatibilities (i.e. Dobzhansky-Muller incompatibilities – DMIs) inherent to the formation of admixed genomes (Runemark et al., 2018b; Trier, Hermansen, Sætre, & Bailey, 2014). Genetic incompatibilities may constrain hybrid lineages long after hybridization has occurred affecting their evolutionary potential (Eroukhmanoff et al., 2017; Runemark et al., 2018b). For instance, selection against DMIs can reduce the availability of variation responsive to adaptive evolution and hence, population divergence and the potential for local adaptation can also be reduced (Runemark, et al., 2018b). Accordingly, any subsequent population divergence within a hybrid species might be largely restricted to loci where the parental species are not fixed for alternative alleles or rely on new mutations occurring after HHS. Thus, the process of HHS includes both the sorting of incompatibilities and fixation of favorable genetic combinations to generate viable and functional genomes. Yet, only few studies have addressed incompatibilities in hybrid genomes (Rieseberg et al., 2003; Runemark, et al., 2018b). Little is still known on how admixture may ultimately constrain or facilitate adaptation in hybrid lineages and how genetic variation in hybrid species is generated and made accessible to selection.

In addition to constraints inherent to admixed genomes, hybrid lineages experience the same challenges as non-hybrid species do, such as intraspecific gene flow constraining local adaptation. Many studies have shown that without geographic isolation, adaptive population divergence can be hampered by gene flow (Hendry & Taylor, 2004; Räsänen & Hendry, 2008; Stuart et al., 2017). Genetic and phenotypic population divergence may therefore be obstructed by gene flow. However, the opposite process can also occur, local adaptation may constrain gene flow, favoring divergence between populations (Gosden, Waller & Svensson, 2015; Räsänen & Hendry, 2008). The examination of factors that may mediate population differentiation (i.e. environmental variation or geography) in conjunction with inference regarding the role of drift and selection is therefore crucial to understand population divergence (Prunier, Colyn, Legendre, Nimon, & Flamand, 2015; Seeholzer & Brumfield, 2018; Wang, 2013). In the specific case of hybrid lineages, it has been argued that incompatibilities could also reduce gene flow (Bierne, Welch, Loire, Bonhomme, & David, 2011), especially when genes under ecological selection are coupled with DMI loci (Seehausen, 2013) which could in turn facilitate local adaptation (Eroukhmanoff, Hermansen, Bailey, Sæther & Sætre, 2013; Trier et al., 2014). Hence, it is important to assess whether local adaptation is more or less constrained in hybrid lineages, compared to non-hybrid taxa, given their specific genomic architecture.

In this study we investigate how the homoploid hybrid Italian sparrow (*Passer italiae*) has evolved since its formation. How constraints and novel genetic variation, linked to admixture, have impacted its genomic evolvability, limiting or favoring its adaptive potential and ultimately population divergence. The Italian sparrow is a homoploid hybrid species resulting from past hybridization between the house sparrow (*Passer domesticus*) and the Spanish sparrow (*Passer hispaniolensis*) (Hermansen et al., 2014; Trier et al., 2014). This hybridization event likely occurred when the house sparrow spread into Europe alongside agriculture, approximately 6 Kyr BP (Hermansen et al., 2011; Ravinet et al., 2018). In mainland Italy its genome is completely admixed with a slightly higher contribution from the house sparrow (Elgvin et al., 2017). It is reproductively isolated from its parental species, with strong post-zygotic barriers associated to mito-nuclear and sex-linked incompatibilities (Elgvin et al., 2017; Trier et al., 2014). Phenotypically and genetically, the Italian sparrow is a mosaic form between its parental species (Elgvin et al., 2017; Hermansen et al., 2011).

Patterns of population divergence and local adaptation at the genomic level have not yet been investigated in the Italian sparrow, nor the extent to which genomic constraints might have affected population divergence in this species. We limited our study to mainland populations across the Italian peninsula, using restriction site-associated DNA sequencing (RAD-seq) and the house sparrow genome assembled by Elgvin et al., (2017) as a reference genome. We assessed population divergence across the Italian peninsula and the role of climatic variation on genomic divergence. Our results suggest that genetic divergence within the Italian sparrow is driven by climatic variation. We report patterns of isolation by environment (IBE), temperature apparently being the main driving factor. We identify some outlier loci of adaptive divergence associated with precipitation and beak height variation. Furthermore, loci inferred to be under selection are located close to genes known to be involved in cranio-facial development in other bird species. Finally, to determine the nature of the genomic divergence patterns found in the hybrid species, we examined genomic divergence in its parental species. Our results demonstrate that most loci involved in local adaptation in the hybrid species are not divergent between parental species, whereas loci involved in local adaptation within parent species seem to have previously been under divergent selection between the parental taxa. Overall, genomic divergence and local adaptation seems to be highly polygenic both in the hybrid and parent species, albeit different loci and genomic regions are involved in adaptive intraspecific divergence.

## METHODS

### Study species and sampling

The Italian sparrow is distributed across the Italian peninsula and a few Mediterranean islands. Of its parental species the house sparrow has a wider native distribution extending throughout large parts of Eurasia whereas the Spanish sparrow is located around the Mediterranean Sea and eastwards to Central Asia (Summers-Smith, 1988). We concentrated on the mainland distribution of the Italian sparrow, excluding populations from the Mediterranean islands as they likely represent independent hybridization events (Runemark, et al., 2018b). We also sampled across the distribution of the parental species.

Birds were caught using mist nets. Blood samples were obtained by puncturing the left brachial vein and stored in standard Queen’s lysis buffer. Individuals were released immediately after sampling to minimise stress. All relevant sampling permits were obtained from the regional authorities.

We sampled a total of 131 (68 males and 63 females) Italian sparrows from 8 populations across the Italian peninsula (Fig. 1A, Table S1) during the spring of 2007, 2008 and 2012. These populations are geographically well spread representing most of the continental distribution of the Italian sparrow.

**Figure 1.**
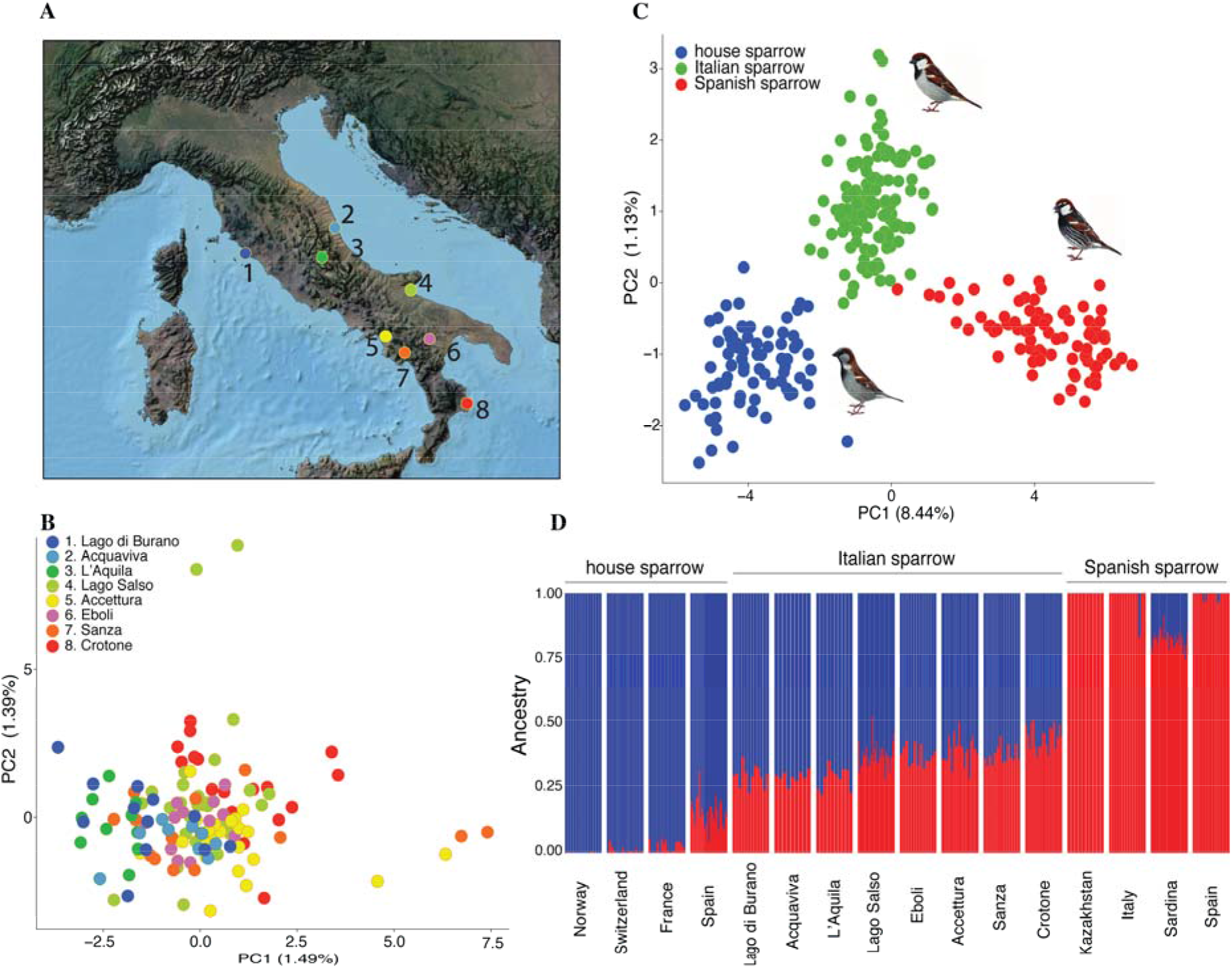
**A.** Map showing the distribution of sampled Italian sparrow populations **B.** Principal component analysis (PCA) to explore genetic divergence within the Italian sparrow (8 Italian populations, 131 individuals and 4387 SNPs). **C.** PCA including the Spanish sparrow (red), house sparrow (blue) and Italian sparrow (green), and **D.** Admixture analysis among the three focal species, based in a vcf file containing 288 individuals (131 Italian, 82 Spanish and 75 House sparrows) and 2737 high-quality SNPs. Localities are ordered following latitudinal distribution.

In addition, we sampled 82 Spanish sparrows (51 males and 31 females) from Spain, Italy, Kazakhstan and Sardinia and house sparrows (n = 75, 49 males, 26 females) from Norway, Switzerland, Spain and France, where locations vary between 13 to 27 individuals sampled (Table S1).

### DNA extraction and sequencing

Genomic DNA was purified from blood samples using Qiagen DNeasy 96 Blood and Tissue Kits (Qiagen N.V., Venlo, The Netherlands) according to the manufacturer’s instructions. The protocol was slightly modified by adding 125 ul of blood stored in Queen’s lysis buffer and warming the elusion buffer (EB) to 40°C to increase yield of DNA. DNA isolates were stored in Qiagen Elution Buffer (EB). Double digestion of the genomic DNA for ddRAD sequencing was performed using EcoRi and MseI restriction enzymes following the protocol by Peterson, Weber, Kay, Fisher& Hoekstra, (2012). Genomic DNA from single individuals was digested and ligated to respective adapters comprising EcoRI and MseI restriction overhangs. Molecular identifier tags were added with PCR amplification. Resulting individual sample libraries were pooled and library pools were size selected for fragments between 500-600bp with gel electrophoresis and extraction of the respective size range. The size selected library pools were then sequenced using Illumina Nextseq500 machine and the 1×75bp sequencing format. On average, 2.4 ×10^6^ single reads were produced per sample. Library preparation, sequencing, demultiplex and trimming of the adapters were performed by Ecogenics GmbH (Balgach, Switzerland) (www.ecogenics.ch).

### Mapping to reference genome and variant calling

RAD sequences were quality checked by FASTQC (https://www.bioinformatics.babraham.ac.uk/projects/fastqc) and mapped to the house sparrow reference genome, assembled by Elgvin et al. (2017), with BWA-MEM (v 0.7.8) (Li & Durbin, 2009) using the default parameters with the exception of using the −M flag allowing Picard compatibility for further analysis. On average 90.1% of reads were mapped across individuals, similar to mapping percentage shown by closely related species (Ravinet et al., 2018). Bam files were sorted by coordinates using Picardtools (v 1.72) SortSam (https://broadinstitute.github.io/picard/). Identification of indels and local realignment was run on the mapped and sorted reads using Genome Analysis Tool kit (GATK)’s RealignerTargetCreator and IndelRealigner (Auwera et al., 2014; Mckenna et al., 2010) with default parameters. We validated bam files with the Picardtools (v 1.72) ValidateSamFile tool.

From the realigned bam file a set of variants were called by GATK (v 3.7) HaplotypeCaller using the following cut off for filtering: a Phred based mapping quality score of 10, soft clipping of the last 5bp without the need to soft clip both ends (-rf OverclippedRead --filter_is_too_short_value and --do_not_require_softclips_both_ends). The resulting individual genomic variant files (gVCF) were then combined by CombineGVCFs and merged using the GenotypeGVCFs tools. As our analyses were based on single nucleotide polymorphisms (SNPs), all indels were excluded using the GATK’s SelectVariants tool. Variants in unplaced scaffolds were removed using the SelectVariants. Individuals with a proportion of missing data greater than 0.75 were discarded.

SNPs were subsequently filtered by quality using Vcftools v. 0.1.14 (Danecek et al., 2011) as follows: proportion of missing data < 0.8, genotype quality > 20, Depth of coverage > 10 and minor allele frequency of 0.02. Finally, non-variant sites present after filtering and discarding missing-data-individuals, were removed using the GATK’s SelectVariants tool with the ‒env parameter.

We ran a PCA and admixture analysis to identify divergence among species, using a VCF file containing the Italian sparrow and its parental species (288 individuals, 131 Italian, 82 Spanish and 75 house sparrows, 2737 high-quality SNPs).

Within-species analyses were conducted using species-specific VCF files by selecting the correspondent samples, merging individual genomic variant files (gVCF) and genotyping using the GenotypeGVCFs and finally recalling variants within species. Filtering was conducted as described above. The Italian sparrow only VCF file contains 131 individuals and 4387 SNPs from 8 populations. VCF files for each parental species were filtered by minor allele frequency of 0.01. A house sparrow only VCF includes 75 individuals across 4 populations and 6503 high-quality-SNPs and a Spanish sparrow VCF file with 1320 SNPs across 82 individuals from Spain, Italy, Kazakhstan and Sardinia.

### Investigating population divergence within the Italian sparrow

To evaluate population structure and divergence in the hybrid species we used a SNP set containing 4387 loci identified across 8 Italian populations (N=131). We ran admixture analysis and principal component analysis (PCA) using the function glPca in the R package ADEGENET 2.0 (Jombart, 2008). We used vcftools v. 0.1.14 (Danecek et al., 2011) and PLINK v. 1.9 (Chang et al., 2015) to transform the vcf file into format files (MAP, RAW, PED and BED) required by ADEGENET.

To assess the potential for isolation by distance in these Italian sparrow populations we used a multiple (and uni-) matrix regression with randomization (MMRR) approach (Prunier et al., 2015; Wang, 2013), correlating geographic distance and genomic divergence (mean pairwise *F*_ST_) across all pairwise comparison of Italian sparrow populations. This method is described in the next section as the same approach was used to identify correlations between genomic divergence and environmental factors and phenotypic traits.

We used Tajima’s D statistics to investigate signals of selection and potentially recent demographic change, which may have occurred post-hybridization. We also calculated mean values of Tajima’s D, nucleotide diversity and *F*_ST_ using vcftools v. 0.1.14 (Danecek et al., 2011).

To identify regions of divergence in the hybrid species, genome scan analyses were performed across the genome for the 8 populations of Italian sparrows. We calculated global windowed F_*S*T_ and nucleotide diversity using a sliding window of 100kb in size with 25-kb steps. We also calculated Tajima’s D on non-overlaping windows of 100 kb given that LD tends to decay within this distance in sparrows (Elgvin et al., 2017).

### Selection, local adaptation and environmental variation

The Italian peninsula varies considerably in climate, thus we investigated whether genomic divergence covaried with environmental variation. Pairwise differences in climatic variables were regressed with the pairwise genetic distance between populations. We analysed five climatic variables obtained from the global climate data server, WorldClim (v. 2.0, http://www.worldclim.org) (Hijmans, Cameron, Parra, Jones & Jarvis, 2005), BIO1 = Annual Mean Temperature, BIO4 = Temperature Seasonality (standard deviation *100), BIO12 = Annual Precipitation and BIO15 = Precipitation Seasonality (Coefficient of Variation). Values were retrieved using the R packages RGDAL (v 1.3-4, Bivand et al., 2017) and SP (v 1.2-4) (Pebesma & Bivand, 2005), with a resolution of 1km. Geographic distance was obtained with the function spDistsN1 from the R package SP (v 1.2-4) and altitudinal data was gathered from the R package RASTER (v 2.6-7) (Hijmans, 2014) and SP (v 1.2-4) using the getData function. We also analysed phenotypic distance in beak traits, mean beak height (BH) and beak length (BL) in each population.

To test for associations between environmental factors, geographic, altitudinal and phenotypic distances and genome-wide divergence we used a uni- and multiple matrix regression with randomization (UMRR and MMRR respectively) approaches (Wang, 2013) and a modification implemented by Prunier et al., (2015), including commonality analysis (CA) to account for multicollinearity (non-independence) among environmental factors. Data was Z-transformed (i. standardization by subtracting the mean and dividing by the standard deviation) to make regression coefficients of the predictor variables comparable (beta weights, Prunier et al., 2015).

MMRR is a multiple regression analysis on distance matrices used to quantify the contribution of environmental and geographic factors to patterns of genetic divergence (Wang, 2013). It allows the quantification of isolation by distance (IBD) and isolation by environment (IBE) and even IBA when a phenotypic variable is included as predictor. One advantage of the method is that it not only resolves whether the dependent and independent variables are correlated but also quantifies the change and directionality (regression coefficients, β_*n*_) that the dependent variable (genomic distance) has with respect to multiple independent variables, i.e. changes in geographic and environmental distances (Wang, 2013). The overall fit of the model is determined by the coefficient of determination (R^2^). Given the non-independent nature of the variables, the significance (*p-values*) of the variable’s effects (β_*n*_) and fit of the model (R^2^) is estimated by randomized permutations of rows and columns of the dependent variable matrix (for a more detailed explanation of the method see Wang, 2013). However, strong multicollinearity among predictors is still a limitation of this approach. Regression coefficients (β_*n*_), fit of the model (R^2^) and their significance can be affected by multicollinearity among explanatory variables (predictors) (Kraha, Turner, Nimon, Zientek & Henson, 2012; Nimon & Reio, 2011; Prunier et al., 2015). To overcome this caveat an incorporation of variance-partitioning procedures via commonality analysis (CA) can be used, which has been implemented by Prunier et al., (2015). This method (CA) developed originally by Newton & Spurrell (1967) decomposes the model coefficients into unique (U) and common (C) variance components (Campbell & Tucker, 1992 in Prunier et al., 2015; K. F. Nimon & Oswald, 2013), allowing to identify the magnitude of collinearity and the unique effect (U) that a predictor variable has on the dependent variable. The common effect (C) represents the proportion of the variance, in the dependent variable, explained by the collinearity of the predictor evaluated and another explanatory variable; while the unique component quantifies the variance explained by the unique effect of the predictor (Prunier et al., 2015).

CA allows determining of the unique (U) and common (C) contributions of each predictor to the response variable (pairwise *F*_ST_) while accounting for collinearity among predictors. The total effect (T = U + C) of each predictor correspond to the total effect that such a predictor has to the variance explained by the model, independently of collinearity with other predictors, and T/R^2^ is the total variation the predictor account for of the variation explained by the model.

These methods have been shown to provide a better resolution to the effects of environment, geographic distance and phenotype, allowing us to identify patterns of IBD, IBE and IBA (Seeholzer & Brumfield, 2018). This approach is ideal for our analysis given the nature of our data. We are interested in understanding whether genomic divergence and gene flow within the Italian sparrow is linked to climatic, geographic and phenotypic variation. We ran univariate and multivariate matrix regression with randomization (UMRR and MMRR, respectively), with 1000 permutations to estimate significance, along with variance partitioning analysis by CA and 95% coefficient intervals of the commonality coefficient were calculated by bootstrapping 1000 replicates, as implemented by Seeholzer & Brumfield (2018).

We used pairwise geographical distance, altitudinal difference, climate disparity per environmental factor and pairwise mean phenotypic distance as predictor matrices and a genomic distance matrix (pairwise *F*_ST_) as the dependent variable. As the number of predictor variables cannot be greater than the number of populations analysed in the MMRR analysis, two models were run evaluating all the predictors.

To identify candidate loci under selection we ran an outlier analysis using Bayescan (v. 2.1 – Foll & Gaggiotti, 2008), for the Italian sparrow and its parental species independently. Bayescan is a Bayesian approach based on the multinomial-Dirichlet model that uses differences in allele frequency to identify candidate loci under selection by decomposing F_ST_ coefficients into population (**β**) and loci (**α**) components; a reversible-jump MCMC evaluates models with and without selection and calculate posterior probabilities of the parameters under the different models (Foll & Gaggiotti, 2008).

We also evaluated how genomic divergence is associated with environmental and phenotypic variation across populations in the hybrid taxon, performing outlier analyses using BayeScEnv, version 1.1 (de Villemereuil & Gaggiotti, 2015), with the same environmental variables ran on MMRR as predictors, including beak height and length. BayeScEnv, as Bayescan, is a genome-scan software based on Bayesian. To account for population structure it uses the F-model and to control for multiple testing, it returns false discovery rate statistics (PEP, q-value). This method allows the incorporation of environmental information as environmental differentiation between populations and the evaluation of associations between allele frequencies divergence and environmental variables.

We ran BayeScEnv using the default parameters, with the 8 populations of the Italian sparrows. As in Bayescan, the parameters **β** used in the neutral model as well as the locus-specific effect using **α** are estimated. However a third model of local adaptation, estimating the parameter ***g***, uses the environmental differentiation information. Significant associated loci were determined by setting a FDR significance threshold of 5% of the correlation q-value of ***g*** (de Villemereuil & Gaggiotti, 2015).

To identify genes associated to local adaptation (coding regions near outlier loci), we used the house sparrow gff annotation file developed by Elgvin et al., (2017). Given that the linkage decay is of approx. 100 kb in the house sparrow (Elgvin et al., 2017) we selected regions at a maximum of 100kb distance from the outlier loci. We calculated Tajima’s D and F_ST_ for windows containing the outlier loci to compare them with the correspondent genome-wide values of those estimates.

### Investigating genomic constraints to population divergence linked to hybridization

To determine the nature of the genomic divergence patterns found in the hybrid species, and how they differ from non-hybrid species, we compared population genomic parameters of the house and the Spanish sparrow to the Italian sparrow.

To identify how highly divergent loci in the hybrid are distributed, for instance whether they are located in genomic regions of high parent species divergence or not, we selected the top 1% loci with the highest *F*_ST_ among Italian sparrow populations and estimated hybrid-parent *F*_ST_ and between-parents (SH) *F*_ST_ values for these same loci. Similarly, we extracted the top 1% loci with the highest *F*_ST_ among house sparrow populations and among Spanish sparrow populations and as for the hybrid species, hybrid-parent *F*_ST_ and between-parents (SH) *F*_ST_ values were estimated for these highly variable loci. We also calculated Tajima’s D for each of the species and compared the observed patterns between species.

To evaluate whether loci involved in population divergence within the Italian sparrow correspond to loci of high or low genetic differentiation between the Italian and Spanish (IS *F*_ST_) sparrows, Italian and house (IH *F*_ST_) sparrows or Spanish and house (SH *F*_ST_) sparrows, we performed logistic regressions on the probability of being a Italian *F*_ST_ outlier. In the models 1% *F*_ST_ outliers are the response variables while additive and interaction effects of pairwise F_ST_ between the three species were tested as predictors.

Evolution of recombination rate variation across the genome may have an effect on patterns of differentiation within and among species (Burri et al., 2015; Ortiz-Barrientos & James, 2017; Ortiz-Barrientos, Engelstädter & Rieseberg, 2016). Thus, we evaluated whether there was a correlation between recombination rate (estimates taken from a linkage map from Elgvin et al (2017)) and genomic differentiation (*F*_ST_) among populations for each of the three focal species (house, Spanish and Italian sparrows) individually.

## RESULTS

### Genomic landscape of population divergence in the Italian sparrow

As found in previous studies (Elgvin et al., 2017; Hermansen et al., 2011) our results support the mosaic nature of the hybrid Italian sparrow genome (Fig. 1C, 1D). To evaluate the genomic variation among populations of the Italian sparrow we performed a PCA and admixture analyses from 8 locations across the Italian peninsula (N=131 individuals, 4387 SNPs. Fig. 1B, S1), covering a wide range of its continental geographic distribution (Fig. 1A). We found no evidence for genome-wide population structure, only moderate among-population clustering.

Estimated parameters of population divergence among Italian sparrows also show a moderate genome-wide population divergence (mean *F*_ST_ = 0.013, π = 3.1813×10^−6^). Nonetheless, it is possible to identify regions of higher divergence in autosomes, with maximun *F*_ST_ values of ~0.17 across populations and high nucleotide diversity (Fig. S2A, S2C).

### Selection, local adaptation and environmental variation

To further understand the genetic differentiation found among populations of the hybrid Italian sparrow we tested patterns of IBD, IBE and IBA using several climatic factors, altitude, geographic and phenotypic distance as predictor variables. We ran UMRR and MMRR models (Wang, 2013) and variance partitioning through commonality analyses (CA) (Prunier et al., 2015; Seeholzer & Brumfield, 2018). We find no evidence for IBD in our dataset (Table 1, Table 2, Table S2). In UMRR (Table 1) geographical distance (GEO) show a non-significant relationship (R^2^ = 0.053, β = 0.004) to genetic differentiation among populations. Its contribution in the multivariate model (MMRR) is non-significant (β = 0.003, *P* = 0.34) and under the commonality analysis the unique (U = 0.03) and common (C = 0.02) effects of variation explained are considerably small (Table 2). Isolation by environment (IBE) seems to be a more determining factor. Results from UMRR and MMRR yield evidence that climate is driving genetic differentiation within the Italian sparrow, suggesting adaptation to climate (or some unmeasured factor correlate of climate). In particular, temperature seasonality explains a significant proportion of the genetic variation, (Table 1, Fig. S3), with a R^2^=0.163 and β weight of β=0.007. The multivariate model including all the climatic factors, altitude and geographic distances as predictors (MMRR – model 1, Table 2), explains 25% of the inter-population variation in F_ST_ within the Italian sparrow (R^2^ = 0.25). Consistent with the results from UMRR, temperature seasonality yields the highest β weight, with a considerable explanatory power (β=0.007) (Table 2), accounting for 8% of the variation explained by the model. However, variance partitioning by CA shows that its contribution is mainly in the common effects (C) with other variables, and its unique contribution is almost negligible (U = 0.003, C = 0.2, Table 2, Fig. S4).

**Table 1.** Univariate regression with randomization (UMRR). 8 populations of the Italian sparrows. Global Pairwise F_ST_ between populations as the response variable. Predictor variables are as following: Annual mean temperature (A.TEMP), Temperature seasonality (TEMP.S), Altitude (ALT), Geographic distance (GEO), Annual mean precipitation (A.PREC), Precipitation seasonality (PREC.S), Beak height (BEAK.H) and Beak length (BEAK.L)

| | $R^2$ | $\beta$ | $t$ | $p$-value |
| --- | --- | --- | --- | --- |
| TEMP.S | 0.163 | 0.007 | 2.251 | 0.048 * |
| A.PREC | 0.061 | -0.004 | -1.302 | 0.260 |
| GEO | 0.053 | 0.004 | 1.201 | 0.264 |
| BEAK.L | 0.036 | 0.003 | 0.991 | 0.358 |
| BEAK.H | 0.036 | 0.003 | 0.991 | 0.341 |
| PREC.S | 0.031 | 0.003 | 0.910 | 0.422 |
| ALT | 0.020 | -0.002 | -0.732 | 0.513 |
| A.TEMP | 0.001 | -0.001 | -0.172 | 0.864 |

**Table 2.** Multiple matrix regression with randomization (MMRR) and coefficients from Commonality Analysis (CA) – MODEL 1. Unique (U), common (C) and total (T) variance partitioning coefficients of each predictor variable to genomic divergence (Pairwise FST), in parentheses the per cent contribution of the predictor to the total variance explained by the model (100 * partition coefficient (U, C or T) / R2). Global Pairwise FST between 8 populations of the Italian sparrow as the response variable. Predictor variables are the following: Annual mean temperature (A.TEMP), Temperature seasonality (TEMP.S), Altitude (ALT), Geographic distance (GEO), Annual mean precipitation (A.PREC), Precipitation seasonality (PREC.S).

**$R^2 = 0.25$**
| Predictor | $\beta$ | $t$ | $p\text{-value}$ | Unique (U) | Common (C) | Total (T) |
| --- | --- | --- | --- | --- | --- | --- |
| GEO | 0.003 | 0.93 | 0.34 | 0.03 (12%) | 0.02 (8%) | 0.05 (20%) |
| A.TEMP | 0.001 | 0.15 | 0.89 | 0.14 (56%) | 0.03 (12%) | 0.16 (64%) |
| A.PREC | -0.004 | -1.16 | 0.32 | 0.05 (20%) | 0.01 (4%) | 0.06 (24%) |
| TEMP.S | 0.007 | 1.96 | 0.05 * | 0.003 (0%) | 0.02 (8%) | 0.02 (8%) |
| PREC.S | -0.003 | -0.60 | 0.57 | 0.001 (0%) | 0.00 (0%) | 0.00 (0%) |
| ALT | -0.002 | -0.29 | 0.78 | 0.01 (4%) | 0.02 (8%) | 0.03 (12%) |

While mean annual temperature explains a considerable amount of the variance (A.TEMP, Table 2) most of it fell into the unique factor (U = 0.14) and its beta weight is non-significant (β = 0.001, *P* = 0.89). Mean annual precipitation shows similar results (A.PREC, T=24%, Table 2). These patterns suggest that there is collinearity between climatic factors. Unique (U) and common (C) contributions to the variation, estimated by CA (Table 2, Fig. S4), show mean annual temperature (T = 0.16) and mean annual precipitation (T = 0.06), as the mayor contributors, accounting for 64% and 24% of the variation explained by the model, respectively (Table 2). However beta weights for these predictors are not significant.

Moreover, when removing mean annual temperature from the model (MMRR – model 2, Table S2) temperature seasonality is no longer significant (*P* = 0.1), supporting the collinearity effect among climatic variables.

Finally, evaluating IBA incorporating beak morphology among the predictors (beak height and length), the univariate (UMRR, Table 1) and multivariate (MMRR, Table S2) models show that these phenotypic traits do not explain a significant amount of the genomic divergence among Italian sparrow populations. The univariate models for each of the beak traits present a not significant **R^2^** = 0.036 (*P* > 0.34), and in the multivariate model (MMRR – model 2, Table S2) beta weights are low (β=0.001 for BEAK.H and β=0.002 BEAK.L) and are also not significant.

To determine whether highly divergent genomic regions are associated with environmental factors and to identify potential genes associated to local adaptation to climate we used a genome scan approach implemented by the software BayeScEnv (de Villemereuil & Gaggiotti, 2015). Five loci were found to be under selection when correlated to environmental variables. On chromosome 5 two outlier loci were associated with mean annual precipitation. One of these presents values of global *F*_ST_ = 0.136 and Tajima’s D = −0.833 among Italian sparrow populations. In comparison, divergence between species pairs for this locus were lower (SH *F*_ST_ = 0.065, HI *F*_ST_ = 0.007 and SI *F*_ST_ = 0.096). A locus on chromosome 15 (with values of *F*_ST_ = 0.172 among Italian populations) was found to associate significantly with mean annual precipitation (Fig. 2A) while presenting high, although no-significant, q-values of ***g*** for mean annual temperature and altitude (Fig. S5A and S5C). Once more, divergence between species pairs were low (SH *F*_ST_ = 0.063, HI *F*_ST_ = 0.029 and SI *F*_ST_ = 0.004) for this locus. Similarly, chromosome 3 and 2 contain one outlier locus each (with a Italian F_ST_= 0.050, SH *F*_ST_ = 0, HI F_ST_ = 0 and SI F_ST_ = 0, and Italian F_ST_= 0.084 and Tajima’s D = 0.999, SH *F*_ST_ = 0.021, HI F_ST_ = 0.036, SI F_ST_ = 0, respectively) associated to precipitation seasonality (Fig. 2B). Within the putative regions of selection (i.e. 100Kb around the outlier loci) we identified 8 genes of interest associated to climatic variation (Table S3). Among which the KAT2B gene has been reported to be involved on heart and limb development (Ghosh et al., 2018) and indirectly associated with the circadian clock (Curtis et al., 2004).

**Figure 2.**
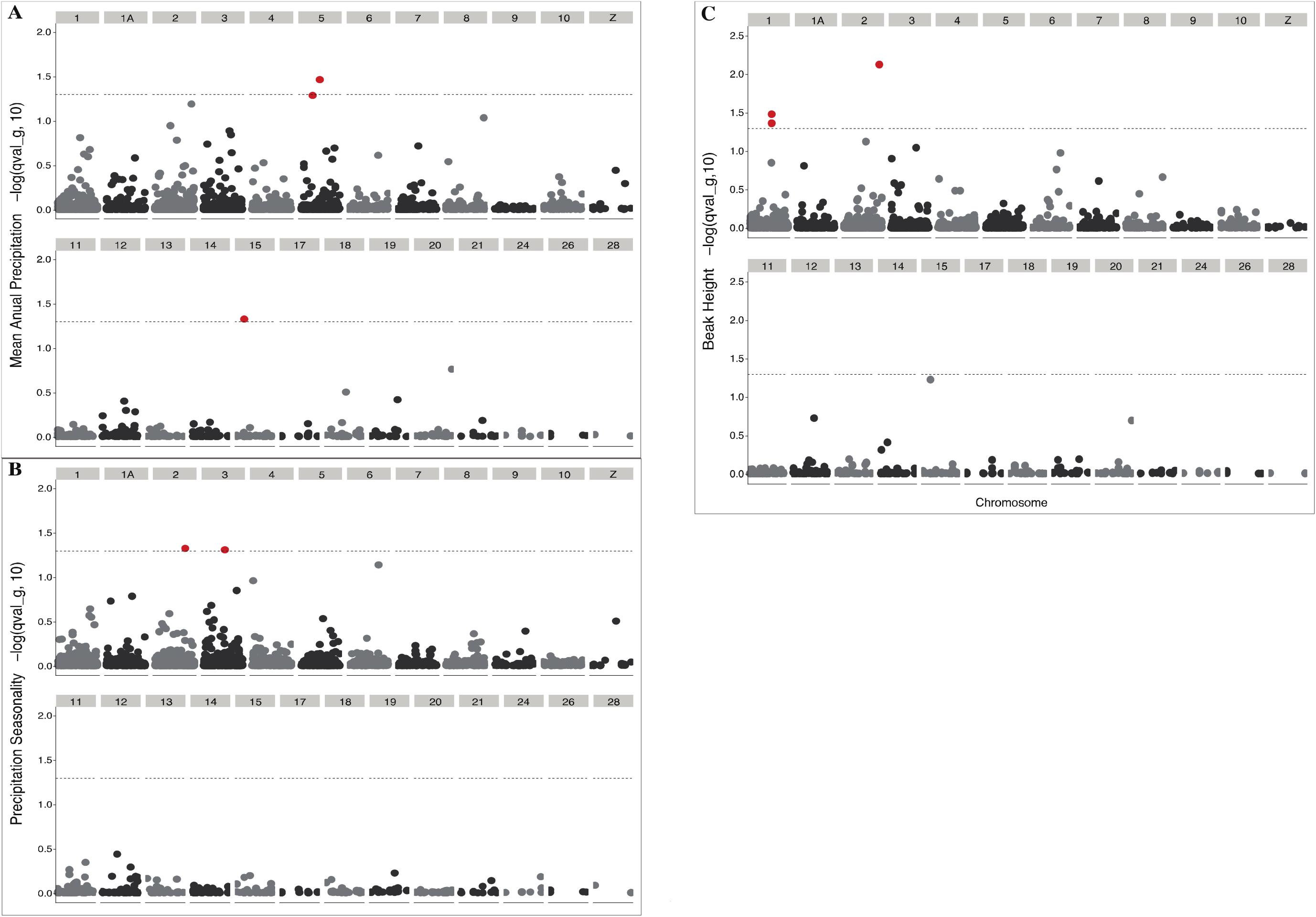
Outlier analysis of local adaptation to climate (BayeScEnv). Manhattan plots of correlation q-values for genetic divergence (SNPs) within the Italian sparrow showing association to climatic factors and a phenotypic trait. Significance level (FDR-corrected) is set at a q-value of < 0.05 (−log10 = 1.3). **A.** Mean Annual Precipitation **B.** Precipitation seasonality and **C.** Beak height.

Using beak morphology traits we identified three candidate loci under selection, two on chromosome 1 and one on chromosome 2, associated with population divergence in beak height (Fig. 2C). No genes were identified adjacent to either of the loci on chromosome 1 while on chromosome 2 we found the genes GJD4 and FZD8. The FZD8 gene is a receptor for Wnt proteins which in turn are involved in patterning the longitudinal embryonic axis and boundary formation at the midbrain-hindbrain junction in vertebrates (Dann, Hsieh, Rattner & Sharma, 2001; Gaudet, Livstone, Lewis & Thomas, 2011).

Further, we used the software Bayescan (Foll & Gaggiotti, 2008) to identify loci under selection across the Italian sparrow populations, independently on whether they are associated to specific environmental factors or phenotypic traits, and compared them with those found by BayeScEnv. We also performed the same analysis in each of the parental species to evaluate whether the hybrid lineage presents similar loci (and associated genes) under selection to those in the parental taxa.

Three outlier loci were identified to be under selection in the Italian sparrow (Fig. 3A). One locus on chromosome 6 and a second locus in chromosome 15, previously identified to be associated to mean annual precipitation by BayeScEnv, however, no genes were found in the vicinity of those outlier loci. Finally, a third locus on chromosome 20 with 6 outlier genes identified. These included genes involved in bone and cartilage development (GDF5) and growth factors (UQCC1) (Fig. 3A, Table S3). The GDF5 gene, also known as BMP-14, encodes a growth differentiation factor protein related to the BMP (bone morphogenetic protein) gene family (Reddi & Reddi, 2009), a gene family involved in skeletal and jaw development (Bleuming et al., 2007; Cerny et al., 2010; Kaucka & Adameyko, 2019). Another of the identified genes (ITCH) is involved in inflammatory signalling pathways and regulate transcription factors involved in immune responses in *Mus musculus* (Shembade, Parvatiyar, Harhaj & Harhaj, 2009; Fang et al., 2002; Gao et al., 2004).

**Figure 3.**
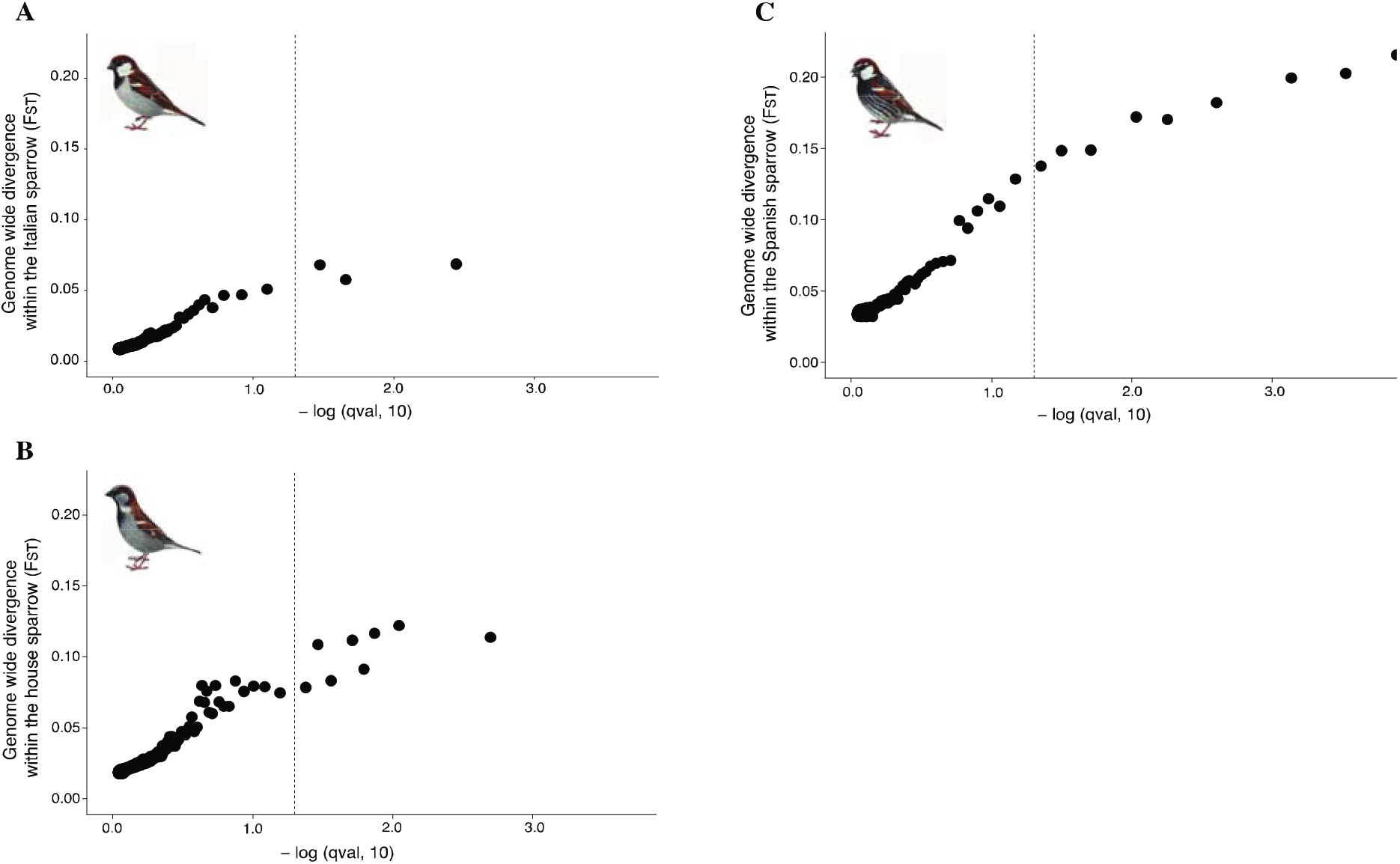
Outlier analysis (BayeScan). Correlation q-values for genetic divergence (SNPs). Significance level (FDR-corrected) is set at a q-value of < 0.05 (−log10 = 1.3). **A.** within the Italian sparrow **B.** within the house sparrow and **C.** within the Spanish sparrow.

In the house sparrow (75 individuals, 6503 SNPs, 4 localities) 8 candidate loci across chromosomes 1, 5 and 8 were identified as being significantly under selection (Fig. 3B, Table S3). 11 genes are found in the vicinity of these candidate loci. The TBX22 gene (T-box transcription factor) is know to be tightly link to the developmental process of palatogenesis in humans. Mutations in this gene causes mouth abnormalities (Braybrook et al., 2002; Braybrook et al., 2001; Marcano et al., 2004).

For the Spanish sparrow (1320 SNPs across 82 individuals from 4 localities) 8 candidate loci were inferred to be under selection with 4 adjacent genes (Fig. 3C, Table S3), including SYNE1 and TDRP which are respectively involved in brain development and male fertility traits including spermatogenesis and sperm mobility (Mao et al., 2016; Zhang et al., 2009).

Only one of the outlier loci was simultaneously identified by both genome scan approaches (Bayescan and BayeScEnv) for the Italian sparrow, suggesting that using a combination of outlier scan analyses gives a better picture of putative regions under selection. Interestingly the hybrid and its parental species seem to be under different selective pressures as none of the outlier loci under selection, nor their associated genes, were found to overlap among the three species.

### Hybrid constraints to population divergence

We compared population genomic parameters between the house, the Spanish and the Italian sparrows to determine whether genomic constrains are playing an important role in the genomic divergence of the hybrid species and to identify differences in genetic variation patterns between the hybrid and non-hybrid, closely related, species.

Population divergence in house sparrow, with maximum values of *F*_ST_ = 0.33 (mean *F*_ST_ = 0.019) and mean nucleotide diversiry of π = 3.219 ×10^−6^ (Fig. S2D, S2F) is similar to that in the Spanish sparrow (mean *F*_ST_ = 0.021, π = 1.923 ×10^−6^), *F*_ST_ up to 0.34 (Fig. S2G, S2I). Divergence in the Italian sparrow is lower than that within each of the parental species, with maximum *F*_ST_ values of ~0.17 (mean *F*_ST_ = 0.013, π = 3.181×10^−6^; Fig. S2A, S2C).

When comparing regions diverging within each of the parental taxa and the hybrid species we found that highly divergent loci (1% *F*_ST_ outliers) in the Italian sparrow are located in regions where different alleles are still segregating in the parental species, i.e, loci with low parental divergence (low SH F_ST_ values). While inherited parental blocks (regions with high values of between-parent-species-divergence (SH F_ST_)) present lower levels of genetic differentiation within the Italian sparrow (Fig. 4A, Fig. S6A). Additionally, none of the highly divergent regions within the hybrid lineage seem to differ substantially from both of the parental species simultaneously, indicative that private alleles do not account for the major population differentiation in the hybrid species (Fig. S6B). Moreover, the probability of being an F_ST_ outlier within the Italian sparrow decreases with the Italian-Spanish (IS F_ST_) genetic divergence (Table 3, *P* = 0.0127). A negative non-significant correlation is also found between the highly divergent regions within the hybrid species and between parental species genetic divergence (SH F_ST_, *P* = 0.0926) (Table 3).

**Table 3.** Logistic Regression on the probability to be an Italian F_ST_ Outlier. Top 1% intraspecific F_ST_ outliers loci selected from a vcf file including the three focal species (131 Italian, 82 Spanish and 75 house sparrows). Outlier loci were identified in a dataset of 2737 shared SNPs between the three species. Italian F_ST_ Outlier loci used as response variable. F_ST_ Outlier threshold=0.06275, Genomic divergence between parental species (Spanish – House (SH F_ST_)) and between the hybrid lineage and each of its parents (Italian – House (IH F_ST_), Italian – Spanish (IS F_ST_)), additive and interaction effects, are used as predictors.

| Model | Predictor | Estimate | Std. Error | p-value |
| --- | --- | --- | --- | --- |
| **Italian $F_{ST}$ Outlier**<br>~ SH $F_{ST}$ | SH $F_{ST}$ | -2.1391 | 1.2719 | 0.0926 |
| ~ IH $F_{ST}$ + IS $F_{ST}$ | IH $F_{ST}$ | 1.9886 | 2.4321 | 0.4135 |
| | IS $F_{ST}$ | -7.6170 | 3.0571 | 0.0127 * |
| ~ IH $F_{ST}$ * IS $F_{ST}$ | IH $F_{ST}$ | 3.2764 | 2.6091 | 0.2092 |
| | IS $F_{ST}$ | -5.9250 | 3.5668 | 0.0967 |
| | IH $F_{ST}$ : IS $F_{ST}$ | -49.4795 | 66.2468 | 0.4551 |

**Figure 4.**
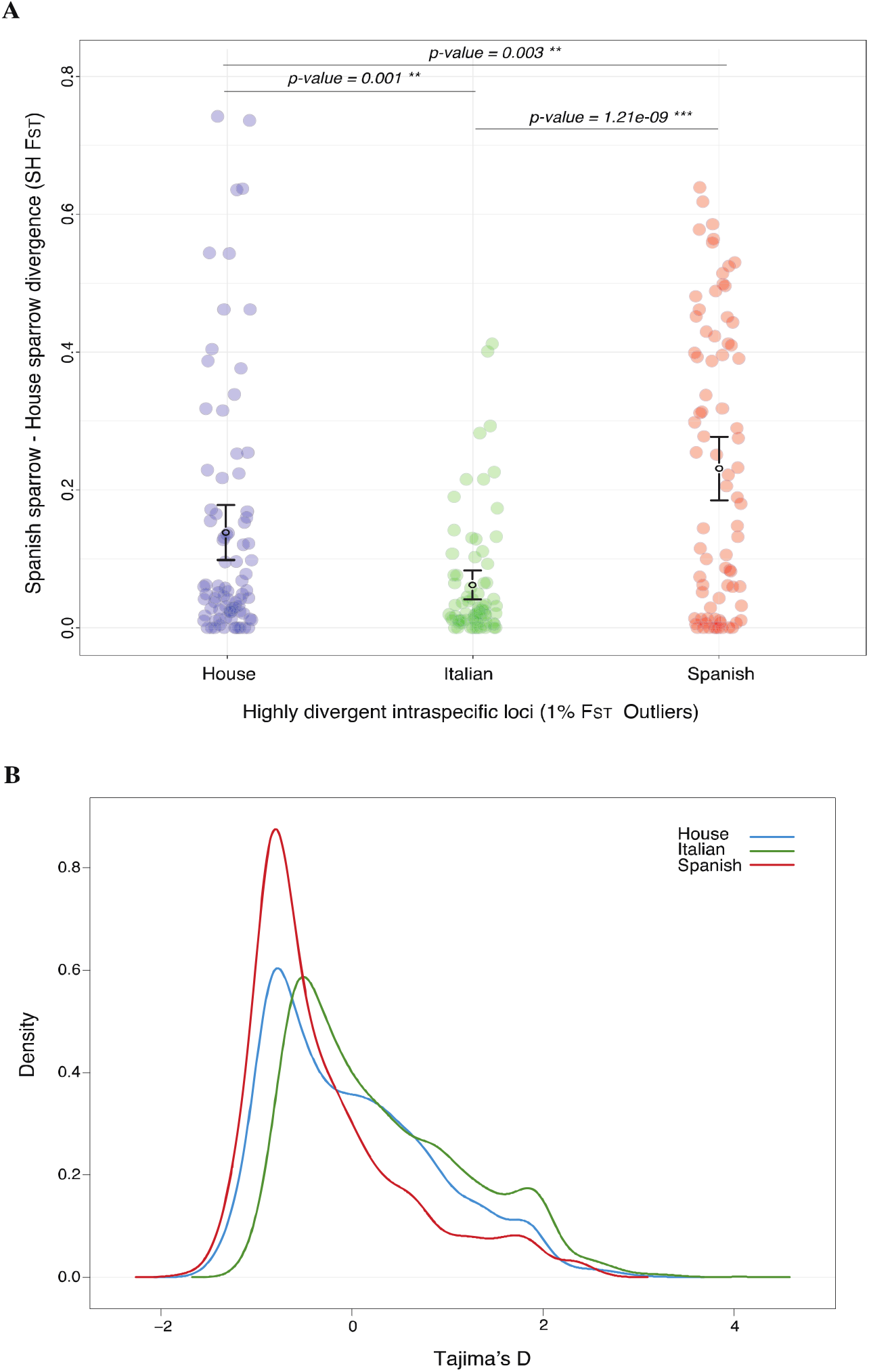
Hybrid constraints to population divergence. **A.** Parental genomic divergence of the intraspecific F_ST_ outlier loci from the three focal species (house sparrow F_ST_ outliers in blue, Italian sparrow in green and Spanish sparrow in red). **B.** Densities of Tajima’s D of the Italian sparrow (green), house sparrow (blue) and Spanish sparrow (red).

In contrast, highly divergent regions within each of the parental species present high parental genomic divergence (high SH F_ST_ values), (Fig 4A)). The 1% outlier loci of within house sparrow *F*_ST_ show higher genomic divergence between the parental species Spanish-House (SH F_ST_) than that within the hybrid species, and the same pattern is found for the Spanish sparrow (Fig 4A). Also, highly divergent loci within each of the parental species do not correspond to those found within the hybrid Italian sparrow (Fig. S6C, S6D).

We did not find evidence supporting that recombination rate could explain the genomic divergence pattern found within the Italian sparrow (R^2^ = −7.986e-06, *P* = 0.607) or within the house sparrow (R^2^ = −1.025e-05, *P* = 0.625). However, for the Spanish sparrow, such a relation is significant (R^2^ = −0.0002629, *P* = 0.0303) (Fig. S7).

Paired samples t-test comparing Tajima’s D between the parental species and Italian sparrows shows that the demographic history and/or selection pressures differ between the species (Fig 4B). We found an overall higher proportion of positive Tajima’s D in the Italian sparrow (t = 15.787, *P* < 2.2e-16, mean = 0.31, Fig. 4B, S2B). However, there are also a number of regions with negative values across the genome (Fig. S2B). The Italian sparrow differ significantly from the house with a mean difference in Tajima’s D of = 0.29 (t = −11.549, *P* < 2.2e-16, Fig. 4B, Fig. S2E) and from the Spanish sparrow where the difference is even larger = 0.58 (t = −16.164, *P* < 2.2e-16, Fig. 4B, Fig. S2H). While the distribution of Tajima’s D in the house sparrow does not deviate from the neutral expectation (t = 0.80993, *P* = 0.418, mean = 0.0132182), the Spanish sparrow presents an overall significantly negative Tajima’s D (t = −9.0721, *P* < 2.2e-16, mean = −0.2735541).

## DISCUSSION

Little is known about how a newly formed hybrid species evolves beyond just a handful of generations. The majority of genomic variation in a hybrid lineage will mainly be derived from admixture. However, novel variation can also be generated from ancient standing genetic variation inherited from the parental species and to a certain extent, through subsequent mutations after hybridization. This variation may ultimately facilitate rapid divergence, whereas genetic incompatibilities may constrain hybrid genomes evolution (Runemark, et al., 2018b), including their potential for local adaptation. Purging of incompatibilities can remove adaptive variation in regions in physical linkage to DMIs (Schumer et al., 2018). In this study we investigated the extent to which populations of a relatively young hybrid lineage have diverged in response to climatic variation and whether their phenotypic traits diverge among populations. We further investigated to what extent divergence in the hybrid occurs at loci where variation is generated by admixture itself, in turn fueling local adaptation.

### Population divergence in the Italian sparrow

We find moderate, but significant genome-wide population divergence, in line with what has been previously found using neutral markers (Eroukhmanoff et al., 2013), and consistent with ongoing gene flow. The young age of this hybrid lineage, thought to be of approx. 6.000 years (Hermansen et al., 2011; Ravinet et al., 2018), may explain this pattern as it may not have been sufficient time for populations to strongly diverge. However, genomic constraints linked to hybridization may also be responsible for this moderate signal of population divergence. Interestingly, this general pattern of differentiation is comparable but somewhat lower than the pattern of population divergence (F_ST_) we report for within each of the parent species, the house sparrow and the Spanish sparrow. Yet, it is difficult to draw further conclusions on the within-species divergence in the parental lineages since populations of the parental species are separated by greater geographic distances than those of the hybrid species, which is likely to affect relative divergence.

Tajima’s D differs between the hybrid and the parental lineages. In the house sparrow, genome wide average of Tajima’s D does not differ from neutral expectation, despite recent work demonstrating a recent population expansion about 6 Kya ago (Ravinet et al., 2018). In contrast, the Spanish sparrow’s average Tajima’s D is negative, while the Italian sparrow present an overall significantly positive Tajima’s D. A pattern of positive of Tajima’s D could be due to nucleotide diversity generated from the hybridization event itself, retained in neutral regions. It would also be consistent with the species having gone through a recent population contraction. Elgvin et al. (2017) reported evidence for balancing selection in some regions of the hybrid genome that would also yield (local) positive Tajima’s D. However, it is diffcult to evaluate to what extent such locally elevated Tajima’s D would influence our genome wide averages. However, we note that a positive Tajima’s D can be consistent with the results we present regarding local adaptation to climate in the hybrid species. One can hypothesise that standing genetic variation could have been maintained by balancing selection and later divergent natural selection led to population differentiation, possibly through the selection of variants playing a role in local adaption to climate (Guerrero & Hahn, 2017). However, it is difficult to conclude and identify the process that could have generated such a positive pattern of Tajima’s D in the hybrid species.

In a previous study, Tajima’s D in the Italian sparrow genome was found to be slightly negative overall (Elgvin et al., 2017) and positive values were mostly located in regions of novel divergence, putatively under balancing selection. The difference between the two studies is puzzling. We tentatively suggest that the difference in the sign of Tajima’s D could either be due the smaller sample size in the previous study that could underestimate the site-frequency-spectrum and to the narrower coverage of the geographic range of the hybrid species in Elgvin et al. (2017), or alternatively, that the type of sequencing (RAD-seq vs. whole-genome resequencing) may matter, as only a portion of the genome is covered by specific restriction sites. Further work is necessary to disentangle the discrepancy between the two studies.

There are several scenarios consistent with the overall moderate population differentiation yet the presence of peaks of divergence across the Italian sparrow genome. Ongoing gene flow among populations can explain the general homogeneity of the genome. Since there is no evidence of a relationship between genomic differentiation and recombination rate, the latter is unlikely to explain the strong peaks of divergence identified. Given the hybrid nature of the Italian sparrow, genomic constraints may be an important factor in its evoultion, hampering population divergence. Consistently, we found loci presenting moderate negative values of Tajima’s D suggesting that some regions in the genome are experiencing purifying selection, potentially linked to purging of incompatibilities. Nonetheless, genetic variation may also have been maintained by balancing selection. Some genomic regions harbour high nucleotide diversity and peaks of divergence among populations, suggesting that there is room for variation in the hybrid genome.

### Selection, local adaptation and environmental variation

Heterogeneity in abiotic factors such as climate and geography can determine patterns of population genomic divergence, either through ecological isolation (isolation by environment IBE, Wang & Bradburd, 2014) where individuals locally adapting to divergent habitats remain separated, facilitating genomic differentiation, or through geographic isolation (Isolation by distance, IBD) where gene flow is limited due to physical distance and geographic barriers (Meirmans, 2012; Slatkin, 1993; Wang, 2013; Wang & Bradburd, 2014).

It has been well documented that in the absence of geographic isolation, gene flow can limit local adaptation, although the directionality of causation of these processes is debatable. Local adaptation can also restrict gene flow enhancing genomic divergence among populations, and even lead to ecological speciation (Gosden et al., 2015; Nosil, 2012; Räsänen & Hendry, 2008). Assessing genomic patterns across a spatially heterogeneous distribution, in correlation with factors that can play a role in genomic divergence, can help us elucidate the processes that have determined population differentiation in hybrid lineages. It can also give insight to the adaptive potential of the species (local adaptation and gene flow reduction) or whether genomic differentiation is essentially a result of genetic drift, where patterns of genetic variation are shaped by low gene flow (Prunier et al., 2015; Seeholzer & Brumfield, 2018; Wang, 2013).

To assess adaptive divergence and gene flow, we evaluated isolation by environment (IBE), isolation by adaptation through beak divergence (IBA) and isolation by distance (IBD). We did not find evidence for IBD or IBA, but the significant correlation between genetic distance and climatic variation is consistent with IBE. Our results suggest that climatic differences, with temperature as the main factor, reduce gene flow between populations in the hybrid Italian sparrow, possibly as a result of local adaptation. Previously, precipitation has been found to be involved in beak morphology variation in the Italian sparrow, and could indirectly be mediating gene flow (Eroukhmanoff et al., 2013). Differential changes in phenotypic traits that respond to specific selective pressure can also contribute to population genomic divergence, thus, in time, local adaptation may sometimes lead to IBA (Edelaar, Alonso, Lagerveld, Senar & Björklund, 2012). However, when directly evaluating beak trait variation as a predictor of genomic differentiation among populations of the Italian sparrow we do not find evidence of IBA.

Patterns of adaptive divergence with ongoing gene flow have also been extensively reported in species of non-hybrid origin (Marques et al., 2016; Martin et al., 2013), which suggest that despite the possibility of constraints reducing the evolvability of this hybrid species (Runemark, et al., 2018b), there is also potential for adaptive divergence leading to local adaptation, as in non-hybrids lineages. In fact, it has been theoretically argued that incompatibilities could facilitate local adaptation by the coupling of genes under ecological selection and DMI loci (Seehausen, 2013). For example, if genomic incompatibilities become trapped in environmentally divergent habitats, coupling with loci involved in local adaptation is possible, which could in turn potentially facilitate diversification within the hybrid lineage (Abbott et al., 2013; Bierne, Gagnaire & David, 2013; Butlin & Smadja, 2018; Seehausen, 2004). This coupling mechanism or divergence hitchhiking (Via & West, 2008), more prone to arise in hybrid lineages around regions of interspecific incompatibilities, could facilitate rapid local adaptation in comparison to other processes of diversifying selection in non-hybrid species (Eroukhmanoff et al., 2013; Seehausen, 2013). To the best of our knowledge, there are no empirical studies that report such linkage between DMIs and regions under natural selection. However, our results and previous studies (Runemark, et al., 2018b) show that genomic constraints play an important role in the formation of the admixed Italian sparrow genome. Moreover, steep genetic clines of putative incompatibility genes involved in reproductive isolation have been found in this hybrid species (Trier et al., 2014). These and our results of genomic divergence in a hybrid lineage may further enlighten the DMI-coupling hypothesis, as a driver of divergent selection, if they are used to evaluate linkage with loci known to be harboring incompatibilities, along with an additional role for phenotypic divergence and local adaptation, in shaping intra-specific gene flow (Eroukhmanoff et al., 2013). However a thorough investigation of such mechanisms at the genomic level is needed to confirm such a hypothesis.

We present for the first time direct evidence on the role of environment in mediating gene flow in a hybrid species, a phenomenon well described in non-hybrid species (Wang & Bradburd, 2014). We also report a few loci where high levels of adaptive genetic differentiation has occurred, some of which are covarying directly with climate variation, suggesting that they are situated in genomic regions linked to local adaptation. Among those loci associated with variation in mean annual precipitation and precipitation seasonality we identified coding genes to be indirectly involved in circadian clock and heart and limb development.

Outlier loci (via genome scans – bayescan) for adaptive divergence between Italian sparrow populations were found to be in the vicinity of genes known to be involved in beak morphology. In particular, GDF5 (growth differentiation factor 5, also known as BMP14 (NCBI)), involved in jaw development in vertebrates (Bleuming et al., 2007; Cerny et al., 2010; Kaucka & Adameyko, 2019) and closely related to the BMP (bone morphogenic protein) gene family (Buxton, Edwards, Archer & Francis-West, 2001; Francis-West et al., 1999a; Francis-West, Philippa, Parish, Lee & Archer, 1999b). This gene family has been found to have a fundamental role in craniofacial development and beak shape and size variation in Darwin’s finches (Abzhanov, Protas, Grant, Grant & Tabin,, 2004; Lamichhaney et al., 2016).

Beak is a trait known to be under strong selective pressure (Lamichhaney et al., 2016; Lamichhaney et al., 2015). Beak size has shown to be a crucial trait underlying the survival of Darwin’s finches after a drought (Lamichhaney et al., 2016) and beak traits in general act as drivers of major evolutionary shifts in Darwin’s finches (Almén et al., 2016; Chaves et al., 2016; Lamichhaney et al., 2016; Lamichhaney et al., 2015). Beak shape variation has been found to respond to environmental divergence affecting food availability in the medium ground finch (*Geospiza fortis*) (Grant & Grant, 2003; Grant & Grant, 2014). Thus, climatic factors could be considered a reasonable proxy for food availability in sparrows (Runemark, Fernández, Eroukhmanoff, & Sætre, 2018). And, it is possible that divergence within genes associated with beak morphology may reflect an adaptive response to variation in food resources found in environmentally different habitats. We also identify two genes linked to population divergence in beak morphology (via BayeScEnv using beak height), among which the gene FZD8, found to be involved in patterning the longitudinal embryonic axis and brain in vertebrates. However, further analyses need to be conducted in order to determine the true underlying mechanisms of divergence between population both at the genetic and phenotypic level.

### Hybrid constraints to population divergence

Evaluating patterns of divergence in the hybrid genome can provide important insights on whether population differentiation is favoured by novel genetic variation or hampered by genomic constraints linked to hybrid incompatibilities. Genomic variation within a hybrid lineage can be generated by novel genetic combinations through rearrangement of parental blocks or polymorphisms generated by the integration of loci that are divergent between the parental species. In these cases we would expect highly divergent loci within the hybrid taxa to be located in regions where the parental species have diverged strongly. Moreover, new mutations occurring after HHS (private alleles) or standing genetic variation inherited from the parental species can also facilitate population differentiation. On the other hand, if genomic constraints linked to admixture are substantially affecting population divergence within the hybrid lineage, by inducing heterozygous genomic regions to rapidly fix one of the two parental alleles, inherited parental genomic blocks should be highly conserved due to incompatibilities.

Our results suggest that genomic constraints and purging of incompatibilities influence population genomic divergence, as loci where parent species are highly divergent seem to remain preferentially fixed for one parental allele across Italian sparrow populations (also evidenced by Runemark et al., 2018b). Furthermore, supporting this constraints hypothesis, we show that genetic variation present in loci that are not differentiated between parental species accounts for most of the genomic divergence found within the hybrid and possibly play a role in local adaptation.

Despite the constrained nature of the hybrid genome, the Italian sparrow has been able to diverge and locally adapt as a response to environmental pressures. One plausible explanation is that variation between populations in the hybrid is generated through inherited standing variation initially present in one or both parent lineages. This variation could be neutral in the parental species, as it seems not to be involved in population divergence in either parent species. Additionally, divergent loci within the hybrid do not differ largely against both of the parental species simultaneously; suggesting that genomic variation within the Italian sparrow is not in areas of novel variation developed privately in the Italian sparrow. However, in depth analyses are needed in order to determine the underlying source of the genomic variation involved in population structuring, and further adaptation, of this hybrid lineage. These results provide a new perspective on how hybridization may impact adaptive evolution, more specifically on how novel genomic variation evolve and is utilized in a hybrid lineage post hybrid speciation, not only through genomic rearrangements linked to admixture and incompatibilities.

Interestingly, patterns of population divergence within the hybrid taxa and each of its parental species seem to differ both qualitatively and quantitatively, suggesting that the admixed nature of the hybrid may somewhat be restricted when compared to species of a non-hybrid origin. Within species genomic variation in the parental lineages (the house and Spanish sparrows) is located mainly in regions of parental divergence while the hybrid species seems to limit its variation to loci that are still segregating in the parental species either because of neutrality in parents or due to constraints limiting the variation in the hybrid. Additionally, there is no overlap of outlier loci under selection, nor their associated genes, among the three species, suggesting differential selective pressures may be operating in addition to specific genomic constraints in the hybrid species. Different population genetic parameters, such as Tajima’s D and nucleotide diversity, also exhibit different patterns, which suggest that the species demographic and evolutionary history may differ and influence genetic patterns across populations. An important factor to be considered in admixed species is the inheritance of traces of different evolutionary histories as well as the private evolutionary path that the hybrid has taken since its formation. Thus, differences in evolutionary patterns among the species are explained by a combination of past evolutionary histories and on-going selective pressures and drift.

## CONCLUSION

Genetic variation within the Italian sparrow appears to be driven by climatic variation, temperature being the main factor; we find evidence for isolation by environment (IBE), which could facilitate ongoing local adaptation. We also find evidence for selection linked to precipitation and phenotypic variation in beak shape, driving population divergence in a polygenic way. Our study supports previous findings suggesting that admixed genomes can be constrained but that nonetheless local adaptation can occur, even under gene flow. Constraints in the hybrid species seem to affect mostly loci of high parental divergence. Adaptive genetic divergence in the hybrid is mainly found in loci that are not differentiated in the parental species and hence less prone to be incompatible in the hybrid. This suggests that purging of incompatibilities is an important element in the evolution of this species. Standing genetic variation inherited from its parent species is a likely explanation for much of the genomic variation in the hybrid species, and some of the variation may be involved in subsequent local adaptation. In contrast, we find little or no evidence that novel variation (private alleles – new mutations occurring after HHS) is important in local adaptation. Coupling of incompatibilities and loci under natural selection could also have facilitated the rapid genomic divergence observed in the Italian sparrow and its effect on gene flow. However, studies addressing these hypotheses directly are necessary to assess causality.

## ACKNOWLEDGEMENTS

We thank S. Vitulano, M. Caldarella, and M. Griggio for assistance in the field. Jo S. Hermansen, Tore O. Elgvin, Stein A. Sæther, Anna Runemark and Cassandra N. Trier for help obtaining the data. Camilla Lo Cascio Sætre and Caroline Øien Guldvog for help with lab work. This work was done in collaboration with the Centre for Naturalistic Studies ONLUS (CSN, Italy) and ISPRA (Italy). This work was funded by The Research Council of Norway grant 297137, the Swedish Research Council and the European Union Marie Sklodowska Curie Action 2011-302504. The Faculty of Mathematics and Natural Sciences (University of Oslo) and Molecular Life Science (MLS) at the University of Oslo.

## DATA ACCESSIBILITY

Genomic data produced in this study will be deposit at the NCBI Sequence Read Archive or at the European Nucleotide Archive (ENA) at http://doi.org/[doi], reference number [reference number]. Other data will be deposited in the Dryad Digital Repository at [accession numbers or DOI will be provided once the data is uploaded].

## AUTHORS’ CONTRIBUTIONS

A.C. and F.E. designed the study; A.C., F.E., M.R. analysed the data; A.C. conducted laboratory work; A.C., F.E. and G-P.S. collected field data; A.C. wrote the manuscript. F.E., M.R. and G-P.S. contributed and commented on earlier drafts of the manuscript.

