## Supporting information for "Intraspecific genomic variation and local adaptation in a young hybrid species"

**Table S1. Specimens sampled.** 131 (68 males and 63 females) Italian sparrows individuals from 8 populations across the Italian peninsula during the spring of 2007, 2008 and 2012. In Addition, we sampled Spanish sparrows (*n* = 82, 51 males and 31 females) from 4 localities (Spain, Italy, Kazakhstan and Sardinia) and house sparrows (n = 75, 49 males, 26 females) from 4 localities in Norway, Switzerland, Spain and France.

**(excel document attached)**

**Table S2. Multiple matrix regression with randomization (MMRR) and coefficients from Commonality Analysis (CA) – MODEL 2.** Unique (U), common (C) and total (T) variance partitioning coefficients of each predictor variable to genomic divergence (Pairwise FST), in parentheses the per cent contribution of the predictor to the total variance explained by the model (100 * partition coefficient (U, C or T) / R2). Global Pairwise FST between 8 populations of the Italian sparrow as the response variable. Predictor variables are the following: Temperature seasonality (TEMP.S), Geographic distance (GEO), Annual mean precipitation (A.PREC), Precipitation seasonality (PREC.S), Beak height (BEAK.H) and Beak length (BEAK.L).

**MODEL 2: Fst ~ GEO + A.PREC + TEMP.S + PREC.S + BEAK.H + BEAK.L	R^2^ = 0.26**

| **Predictor** | **β** | ***t*** | ***p-value*** | **Unique (U)** | **Common (C)** | **Total (T)** |
| --- | --- | --- | --- | --- | --- | --- |
| GEO | 0.003 | 0.58 | 0.60 | 0.01 | 0.04 | 0.05 |
| A.PREC | -0.004 | -1.22 | 0.30 | 0.11 | 0.05 | 0.16 |
| TEMP.S | 0.007 | 1.75 | 0.10 | 0.05 | 0.01 | 0.06 |
| PREC.S | -0.002 | -0.42 | 0.72 | 0.01 | 0.02 | 0.03 |
| BEAK.H | 0.001 | 0.20 | 0.89 | 0.00 | 0.04 | 0.04 |
| BEAK.L | 0.002 | 0.46 | 0.62 | 0.01 | 0.02 | 0.03 |

**Table S3. Genes under selection**. Given a 100kb of linkage decay in the house sparrow, genes were identified within regions at a maximum of 100kb distance from the outlier loci identified with Bayescan and BayeScEnv.

**(See excel file attached)**

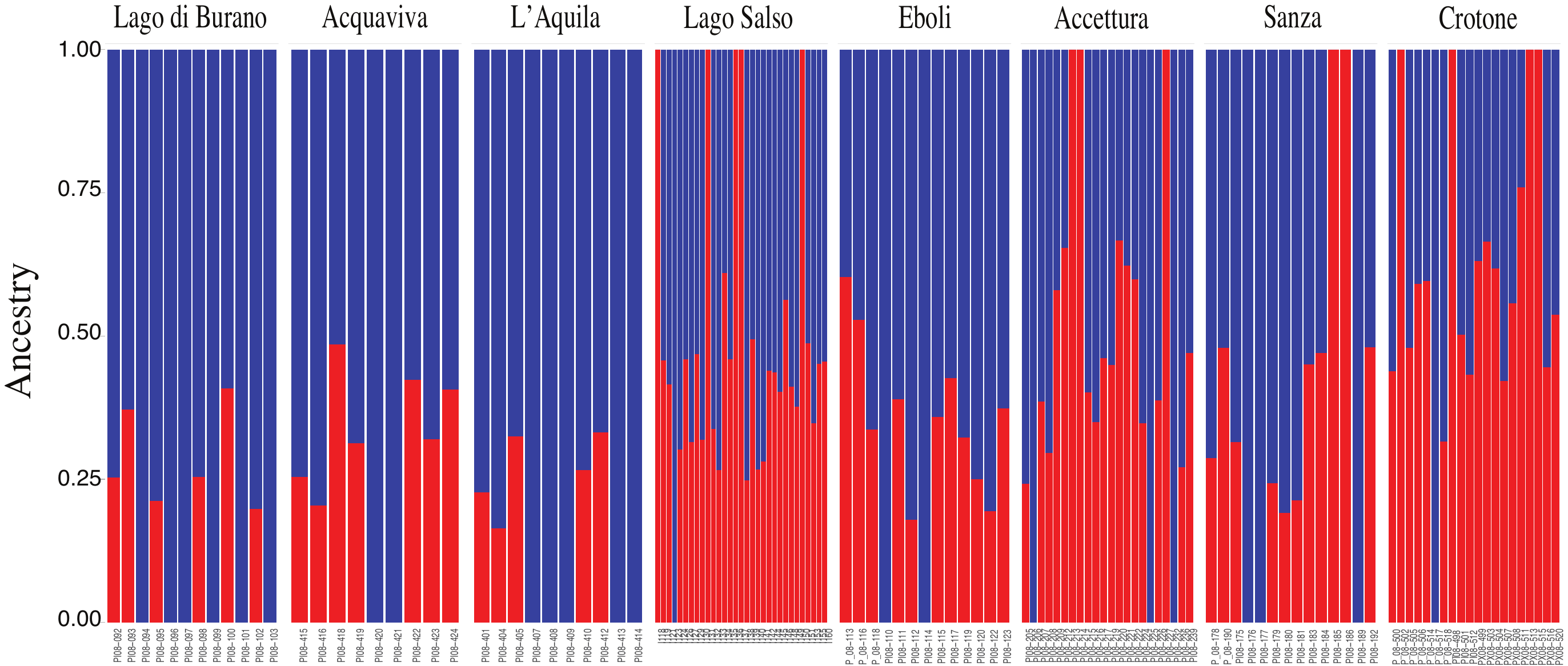

**Figure S1.** Admixture analysis for the Italian populations, 131 samples from 8 populations across the Italian peninsula, 4387 high-quality SNPs.

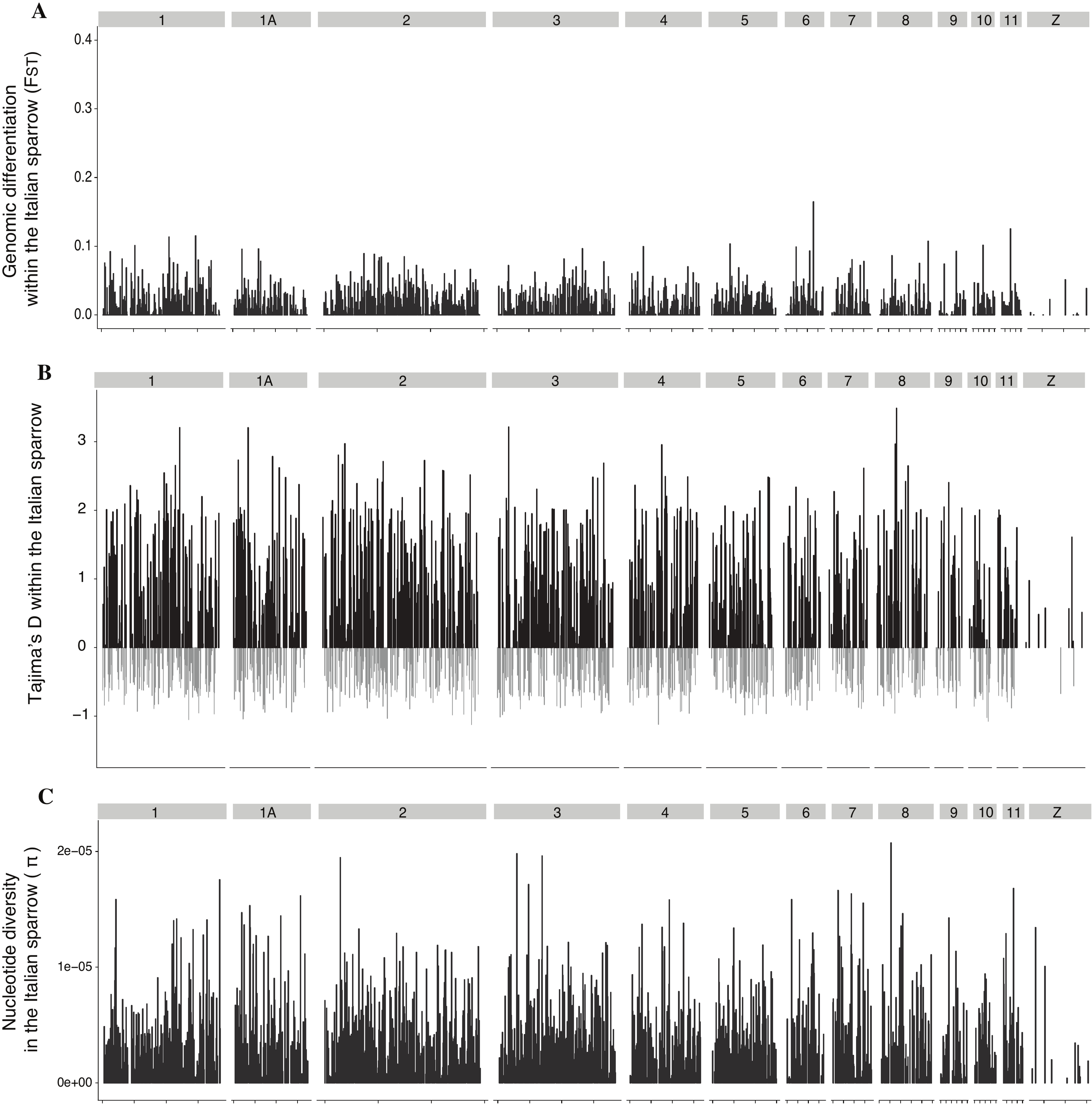

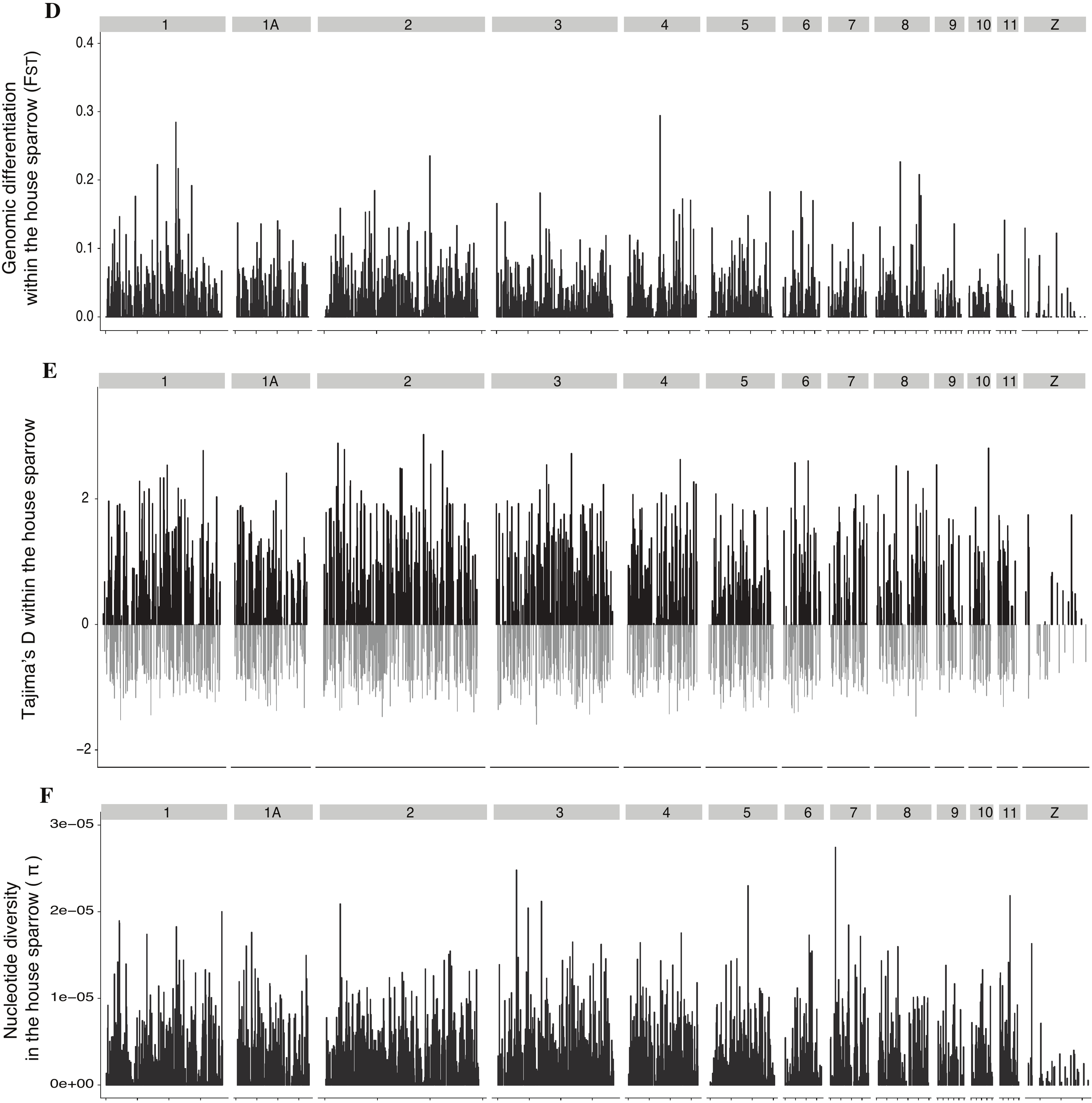

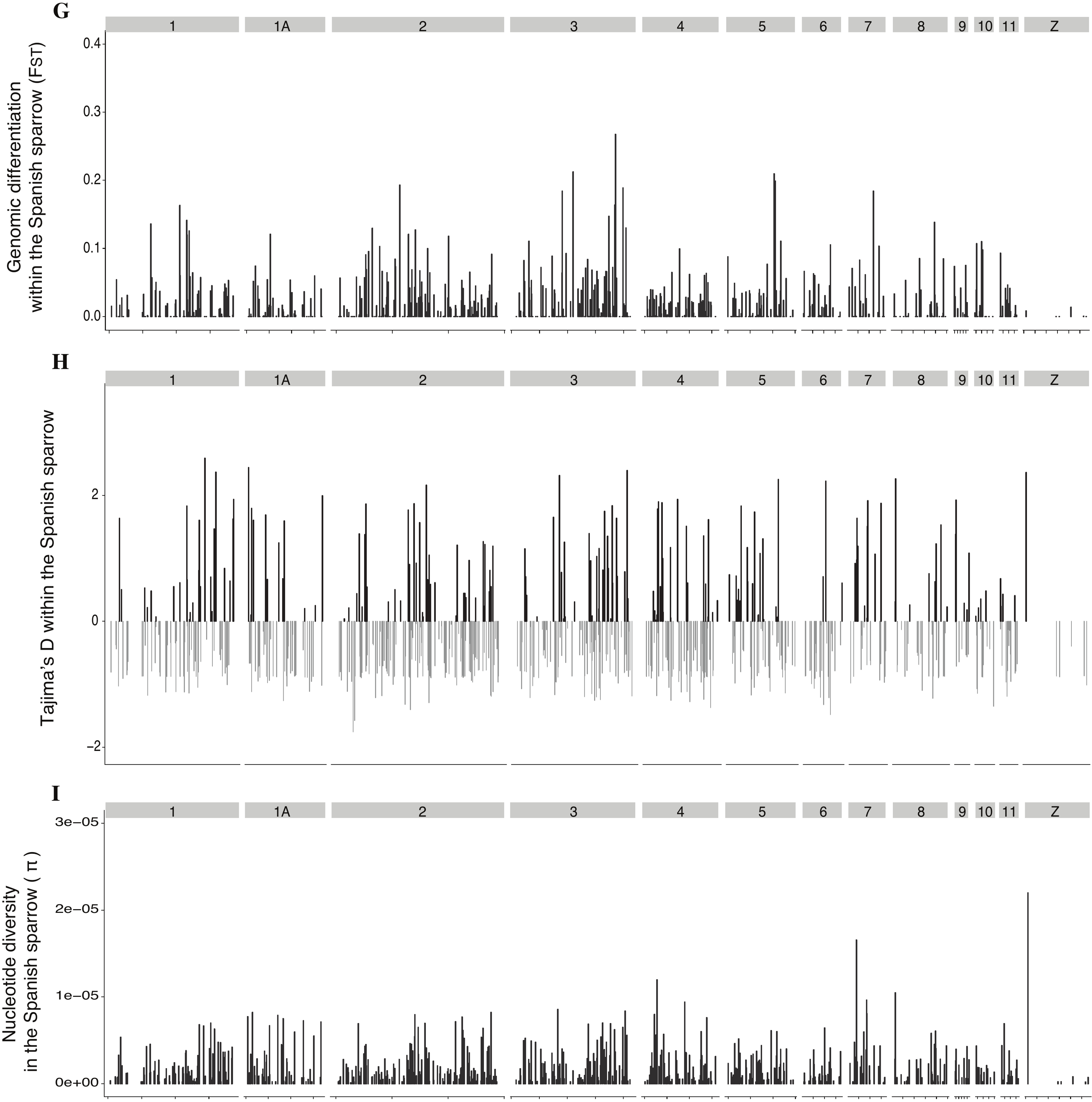

**Figure S2. Genomic landscape of the Italian (A, B, C), house (D, E, F) and Spanish (G, H, I) sparrow.** **A.D.G.** Genome wide population differentiation, global *F*ST estimates for 100kb sliding windows with a window step of 25kb. **B.E.H.** Estimates of selection and demography ­- Tajima’s D - for 100kb non-overlapping windows. **C.F.I.** Nucleotide diversity (π), for 100kb non-overlapping windows with a window step of 25kb. Estimates are based on 4387 SNPs of 131 individuals across 8 populations of the Italian sparrow.

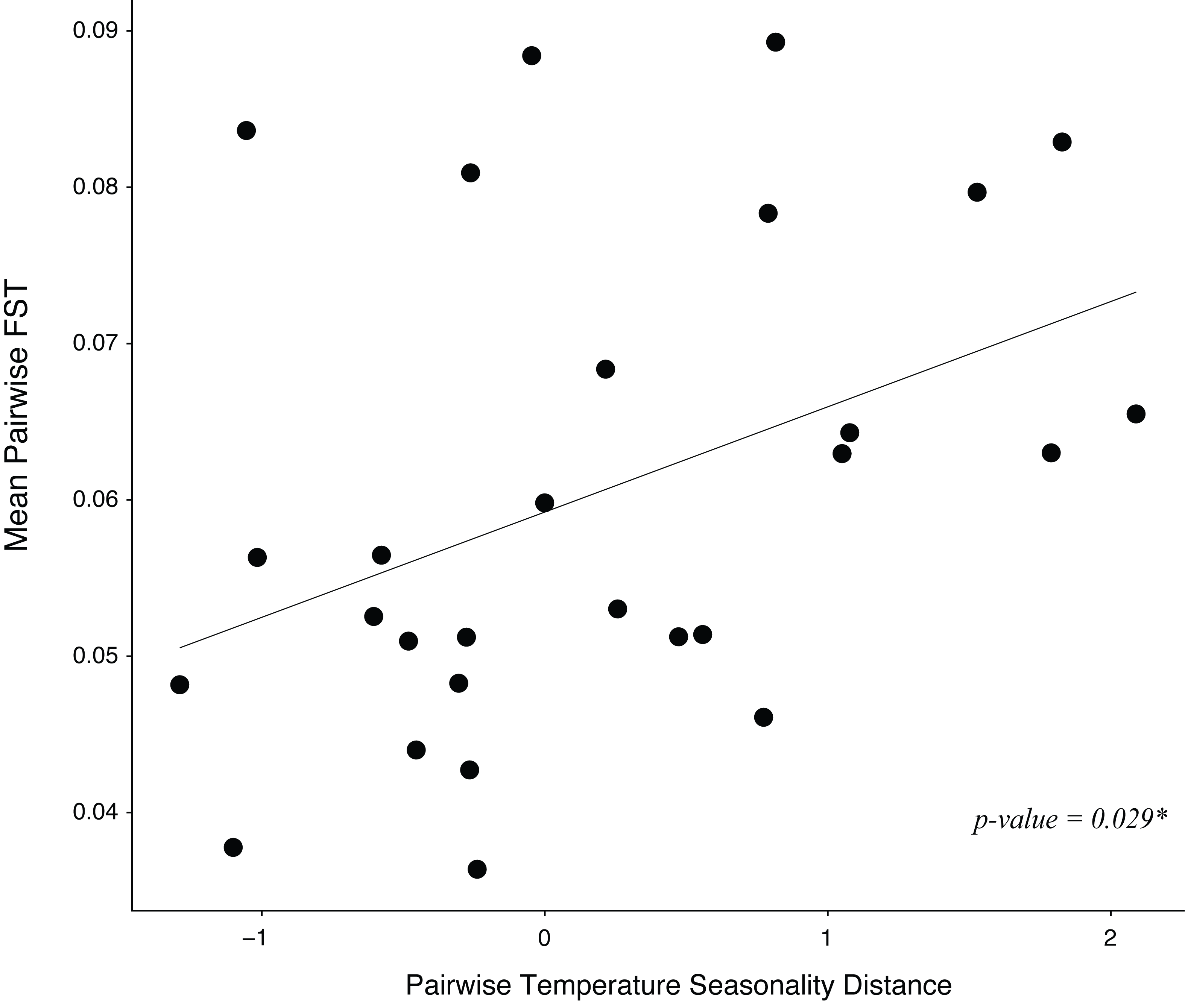

**Figure S3. Local adaptation to climate.** Correlation between temperature seasonality distances and pairwise genomic distance between 8 populations of the Italian sparrow (pairwise FST).

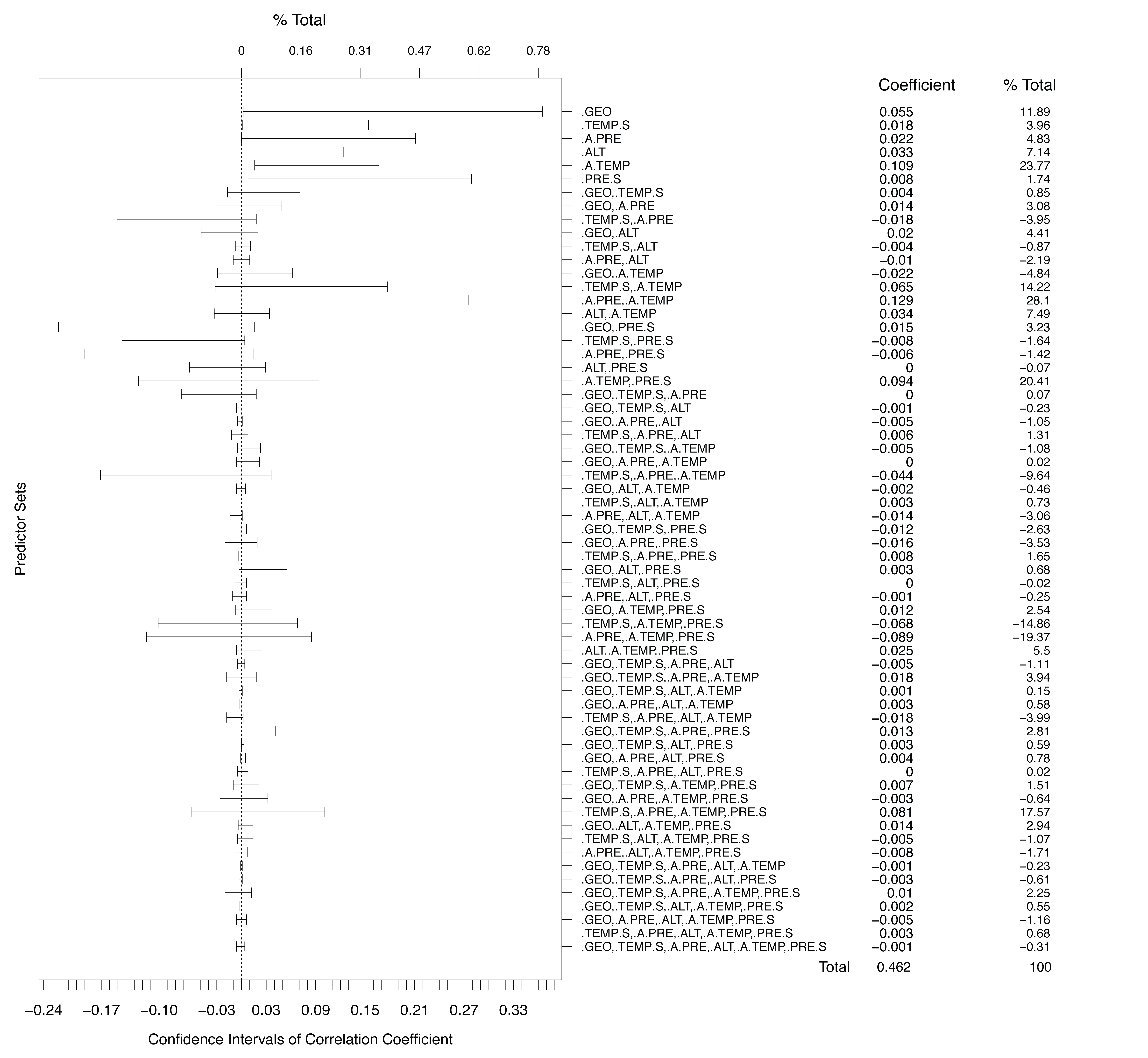

**Figure S4.** Communality analysis (CA) of environmental variables explaining genetic divergence in the Italian sparrow. Communality coefficients and 95% confidence intervals computed using bootstrap with 1000 replicates. Coefficients represent the variance explained by the predictor. *% Total* is the percentage of variance explained, in the variation explained by the model (R^2^), by the predictor, it sum to 100%.

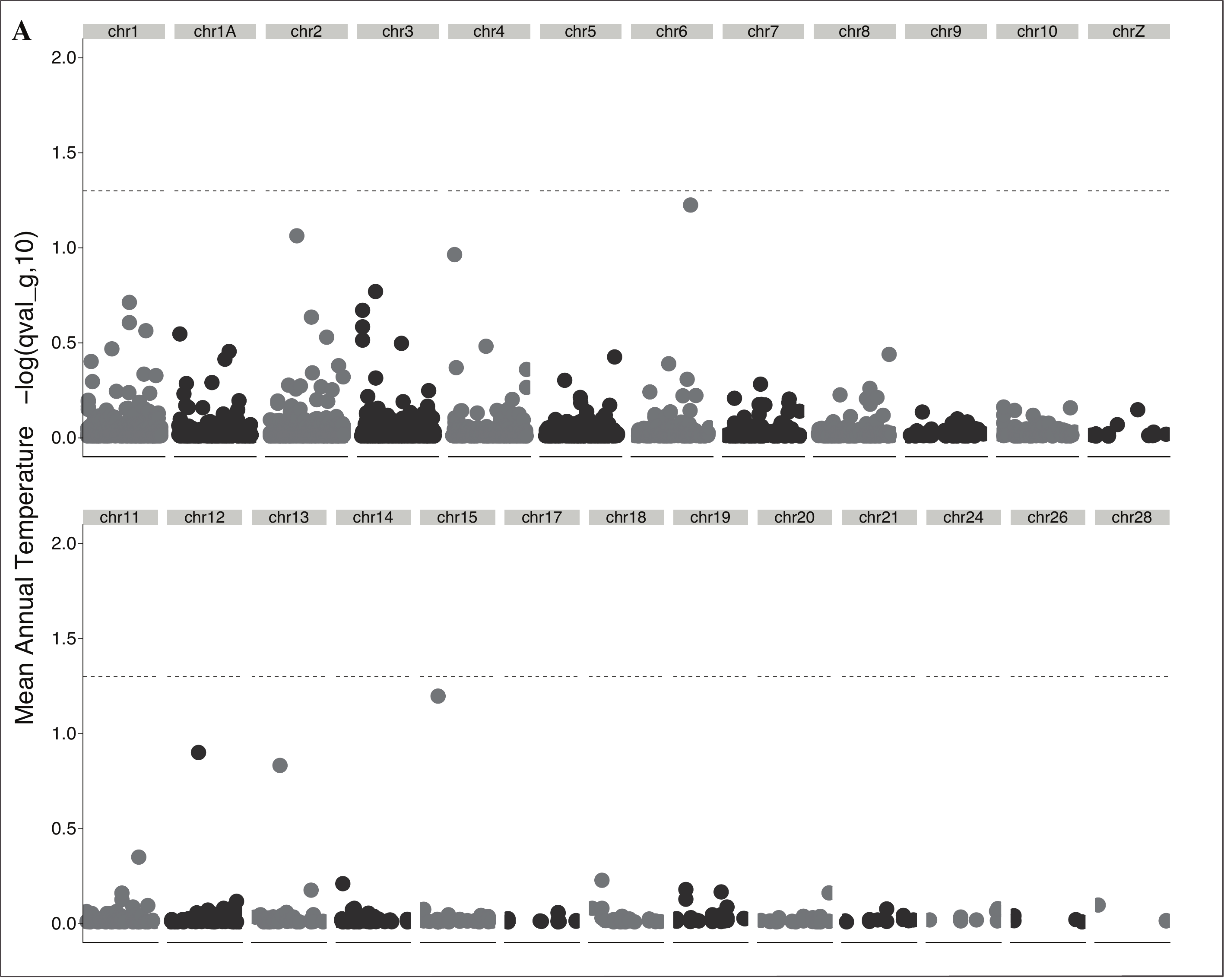

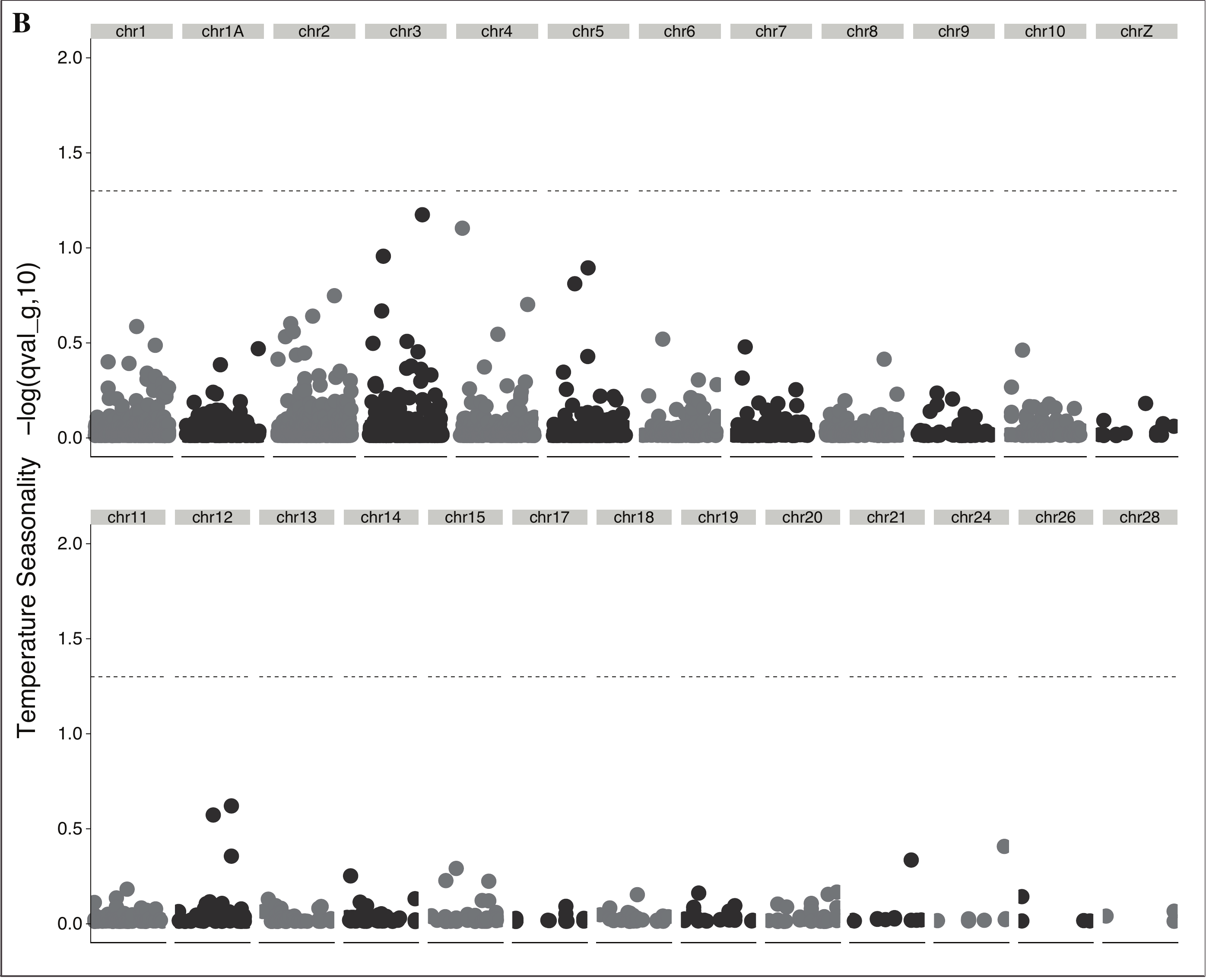

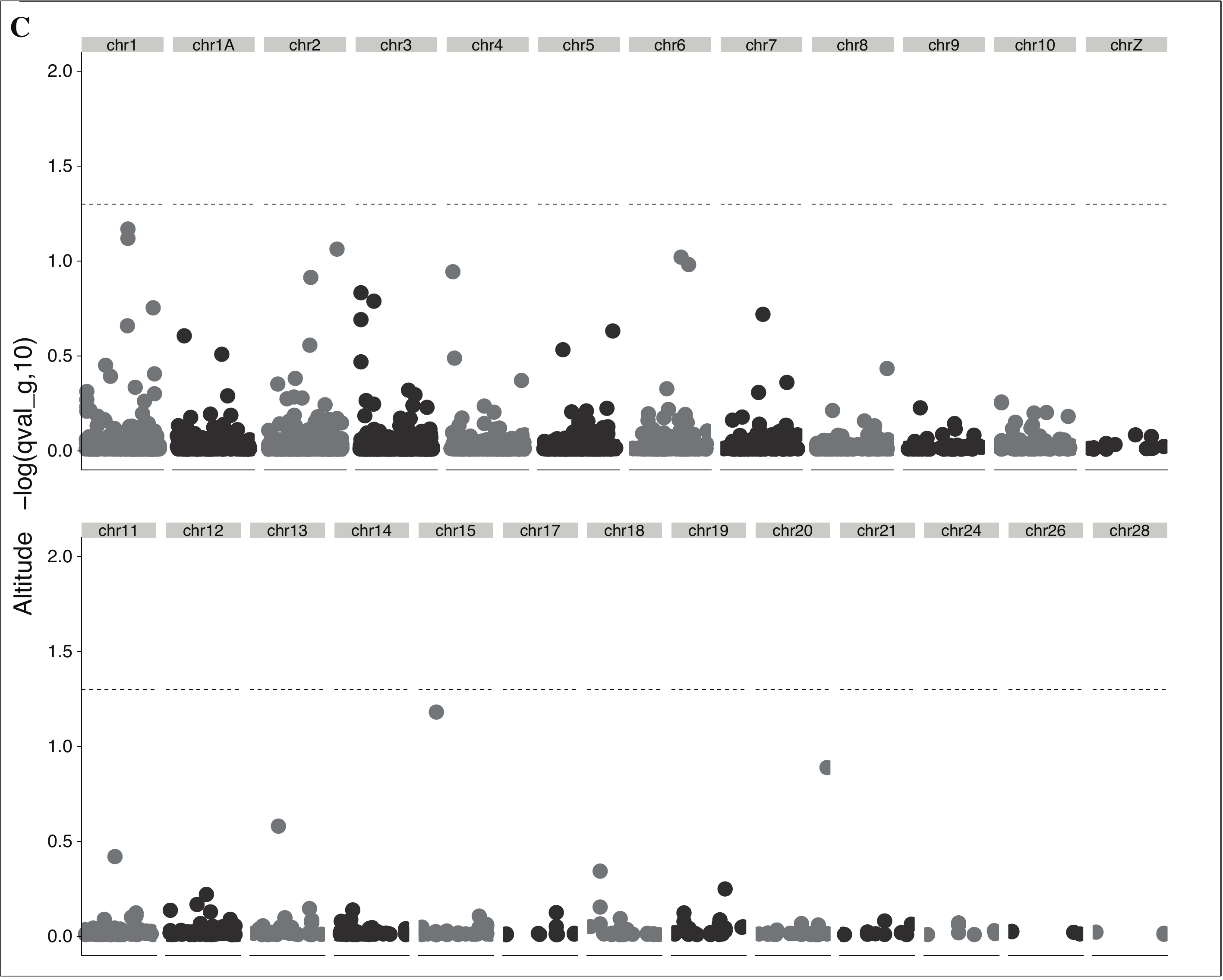

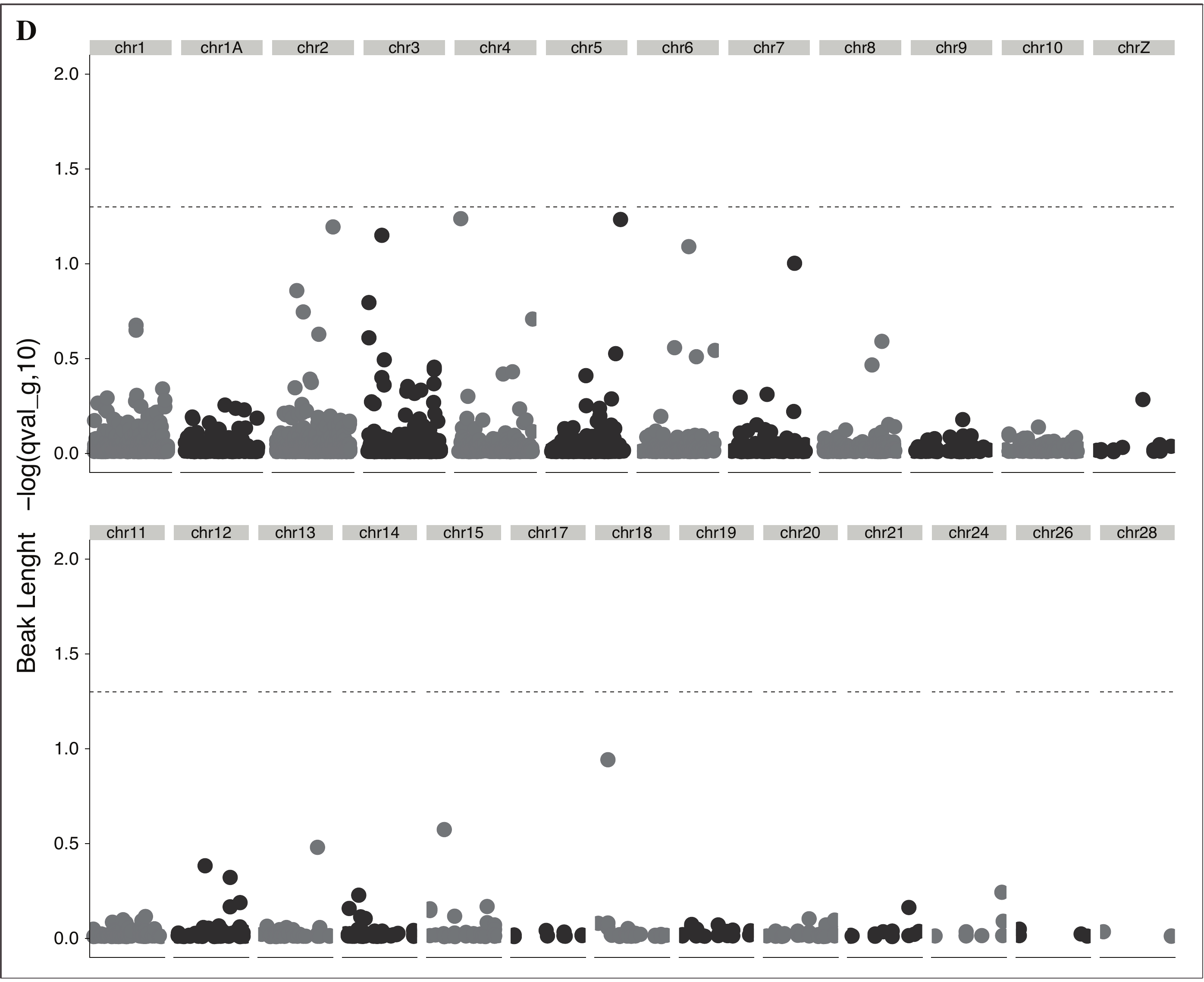

**Figure S5. Outlier analysis of local adaptation to climate and one phenotypic trait (BayeScEnv).** Significance level (F­DR-corrected) was set at a q-value of < 0.05 (-log10 = 1.3). Manhattan plots showing association of Italian genetic divergence (SNPs)_­_­ to **A.** Mean Annual Temperature **B.** Temperature seasonality **C.** Altitude and **D.** Beak length.

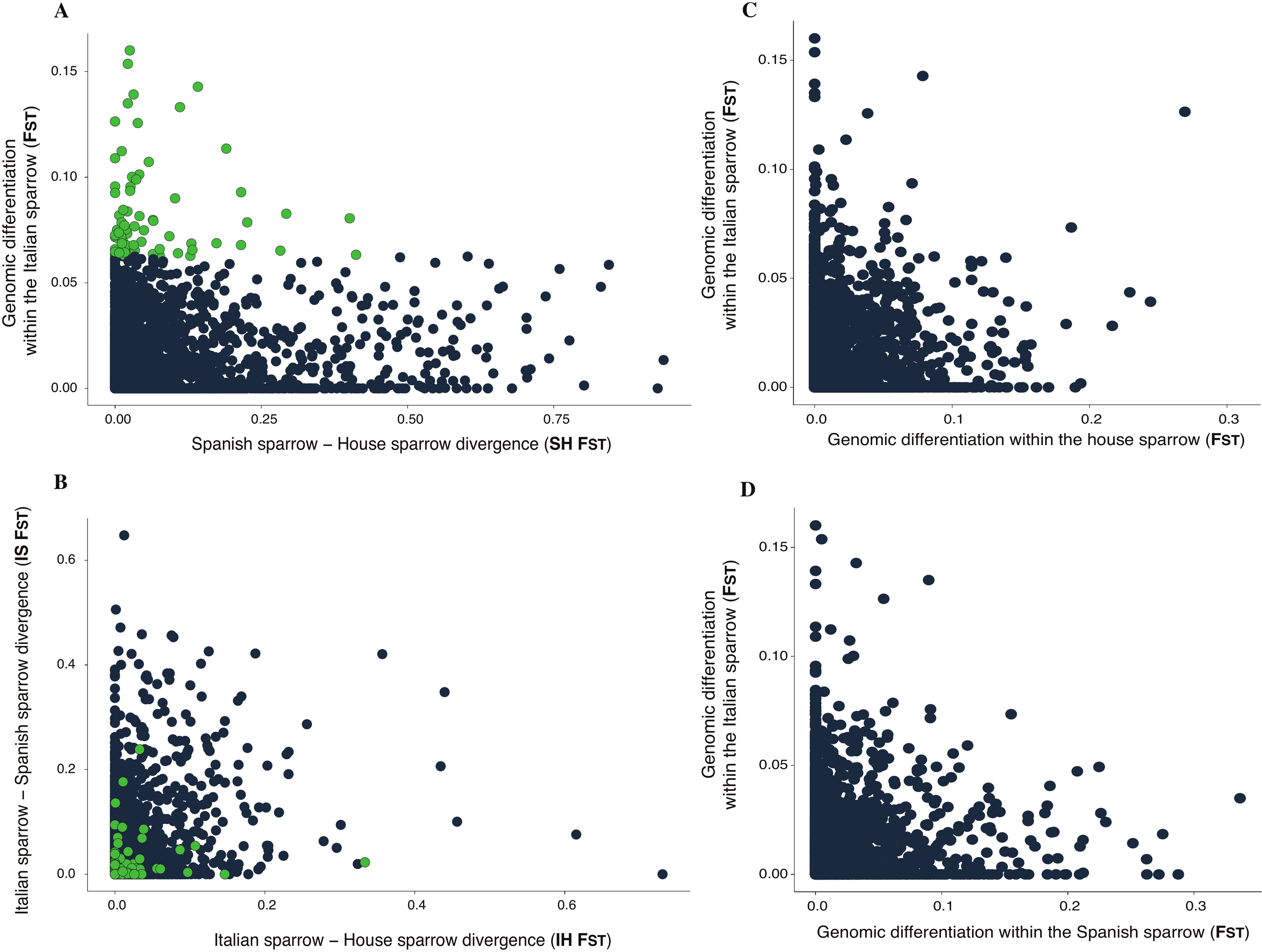

**Figure S6. Hybrid constraints to population divergence. A.** Genomic differentiation within the Italian sparrow (F_ST_) and its parental species (SH F_ST_) **B.** Genomic divergence of the Italian sparrow and each of its parental species (Italian – House sparrow divergence (IH F_ST_) and Italian – Spanish sparrow divergence (IS F_ST_)), with highlighted within Italian sparrow F_ST_ outliers in green. Genomic differentiation within the Italian sparrow v.s. genomic differentiation within each of the parental species, the house **(C.)** and the Spanish **(D.)** sparrows.

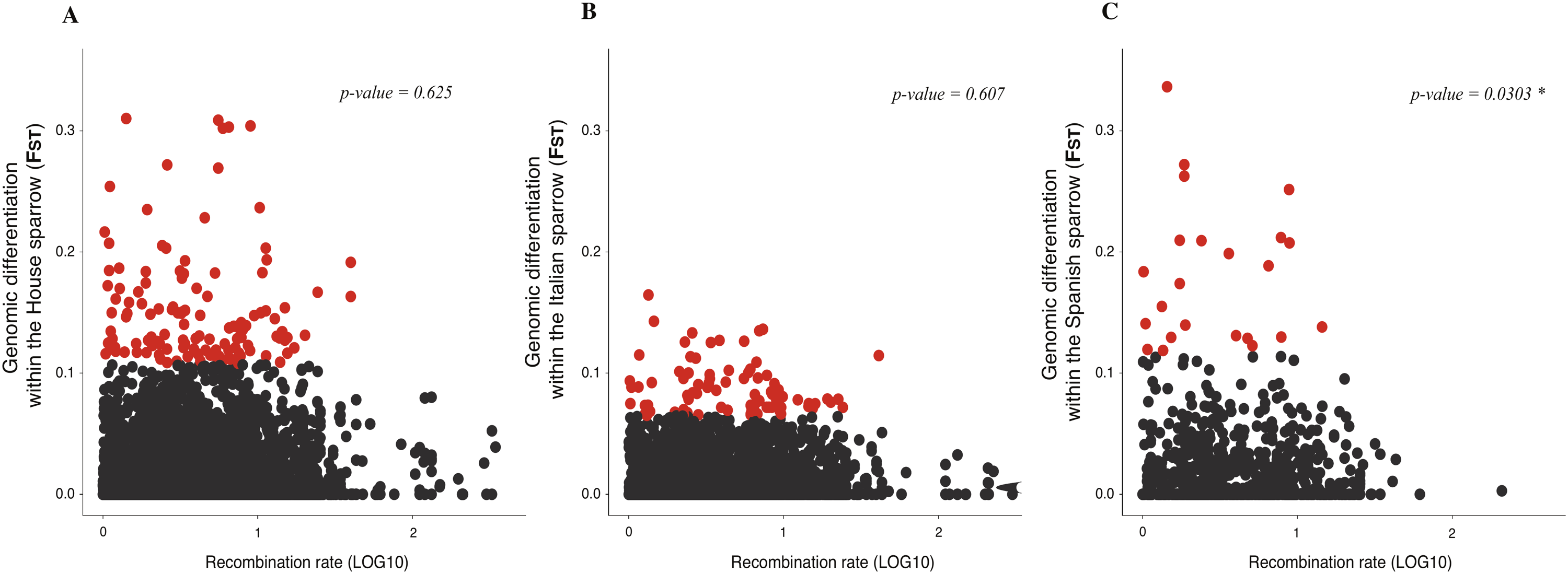

**Figure S7.** Recombination rate v.s. genome wide divergence (F_ST_) **A.** within the house sparrow **B.** the Italian sparrow and **C.** the Spanish sparrow.
